# Discovery of compounds targeting human acute myeloid leukemia stem cells via a novel high-throughput screen

**DOI:** 10.64898/2026.09.09.750169

**Authors:** Isabella A. Iasenza, Safia Safa-Tahar-Henni, Hassan Dakik, Manon Saby, Bahareh Jafari, Patricia Arreba-Tutusaus, Meaghan Boileau, Andrea L. Neumann, Josée Hébert, Sonia Cellot, Frédéric Barabé, Brian T. Wilhelm, Kolja Eppert

## Abstract

Acute myeloid leukemia (AML) is sustained by leukemic stem cells (LSCs), which must be eradicated for durable remission yet remain therapy-resistant. Here, we present a scalable platform to identify compounds that eliminate LSCs by performing the first large high-throughput drug screen directly on human LSC-enriched cells. Using this system, we screened 11 142 compounds and identified 20 inhibitors selective for LSCs. For three candidates, BIO-acetoxime, SJB2-043 and UMxxxxx03, we confirmed anti-LSC activity across multiple primary patient samples and validated efficacy through xenotransplantation assays. Single-cell RNA sequencing uncovered convergent and distinct anti-LSC mechanisms of action, including USP1 inhibition, which suppressed stemness and cell-cycle programs while promoting metabolic and inflammatory stress responses. Collectively, the LSC-enriched fraction was eliminated through apoptosis or differentiation. Together, this work establishes the first platform to directly screen LSCs at scale and identifies novel vulnerabilities and therapeutics capable of eliminating the root of AML.

## Introduction

Acute myeloid leukemia (AML) is an aggressive hematologic malignancy characterized by the clonal expansion of immature myeloid blasts in the bone marrow and blood, ultimately leading to hematopoietic failure (1). Although intensive chemotherapy followed by consolidation achieves remission in approximately 70% of patients, relapse is common and remains the primary cause of mortality (2). A major contributor to treatment failure is the persistence of leukemic stem cells (LSCs), a rare, therapy-resistant population that sustains the disease (3, 4). Thus, the identification of novel therapies that efficiently and selectively target LSCs has the potential to improve long-term responses and survival in AML.

While LSCs share features with normal hematopoietic stem cells (HSCs), they rely on distinct biological programs for survival. These include dependencies on oxidative phosphorylation (OXPHOS), persistently reduced reactive oxygen species (ROS), and increased NF-κB signaling (5, 6). Venetoclax, a BCL-2 inhibitor that targets LSCs by disrupting amino acid– driven OXPHOS, exemplifies the clinical potential of LSC-directed therapy (6–8). In combination with azacitidine, venetoclax has improved outcomes for older or unfit AML patients (9, 10). These findings reinforce both the therapeutic promise and the ongoing need for novel LSC-directed strategies that engage additional biological vulnerabilities.

Despite the appeal of selectively targeting LSCs while sparing HSCs, prior drug discovery screening efforts in AML have failed to translate into the clinic. A central limitation is that these approaches have predominately screened against bulk AML cells or immortalized cell lines (11–19). These approaches lacked a functional readout of LSC activity. As a result, compounds identified through these strategies were found to reduce overall blast burden but failed to eliminate LSCs, driving relapse. Efforts to address this gap using surrogate models or small libraries have remained insufficient, as they do not fully recapitulate the biology of human LSCs or only assess single pathways.

Here, we established optimized conditions to expand and purify LSC-enriched cell populations, enabling direct interrogation of this rare population at scale. Using this platform, we performed the first large scale high-throughput screen (HTS) directly on purified LSCs, testing 11 142 structurally diverse compounds. The screen identified 20 LSC-selective hits and revealed previously unrecognized and targetable LSC dependencies, while establishing a scalable platform for the discovery of therapies that directly eradicate the malignant populations driving AML.

## Materials and Methods

### High-throughput chemical screen and secondary dose-response

OCI-AML-8227 cells were enriched for CD34⁺ cells by magnetic selection and cultured in SFEM II supplemented with cytokines as described in Supplementary Methods. CD34⁺ cells (250 cells/well) were dispensed into 384-well plates containing compounds from a library of 11 142 small molecules. Compounds were tested at 1 μM (commercial and FDA-approved libraries) or 2 μM (investigational and proprietary libraries), with a final DMSO concentration of 0.1%. SR1 (Cat. No. #72342) and UM729 (Cat. No. #72332) (STEMCELL Technologies) were included at 500 nM during primary and secondary screening. For the secondary dose-response screen, 101 compounds were tested in 20 concentrations. After six days, viability was assessed using the CellTiter-Glo® Luminescent Assay (Promega). Primary and secondary screen data were normalized and analyzed as described in the Supplementary Methods.

To assess toxicity toward normal hematopoietic cells, CD34⁺ cells isolated from fresh human cord blood were screened under identical conditions at 5 000 cells/well. All cord blood and patient samples were obtained with informed consent under approved institutional protocols.

### Tertiary screen by flow cytometry

Thirty-three candidate compounds were tested at six concentrations (0.1% DMSO, 2.5, 7.5, 31, 150, 500, and 2 000 nM) using an Echo 555 acoustic dispenser (Labcyte, 0.1 μL/well) in OCI-AML-8227 cells for six days. Cells were then stained with antibodies to CD34 (APC, RRID:AB_1877153), CD38 (PE, RRID:AB_2561900), CD15 (FITC, RRID:AB_314196) (BioLegend), and SYTOX™ Blue Dead Stain (Life Technologies). Flow cytometry was performed on a BD LSRFortessa equipped with a high-throughput sampler (HTS) (BD Biosciences). A threshold LC₅₀ < 500 nM in CD34⁺ cells was used for compound selection.

### Apoptosis assay

OCI-AML-8227, OCI-AML-20, and OP9 cells were treated for three days with seven candidate compounds (2 or 5 μM) or cytarabine as a positive control. Apoptosis was assessed by Annexin V/7-AAD staining and flow cytometry within immunophenotypically defined cell populations. Adult AML patient samples obtained from the Quebec Leukemia Cell Bank (BCLQ) under approved research ethics protocols were cultured in OCI-AML-8227 medium supplemented with SR1 and UM729 and treated under identical conditions. Flow cytometry panels, staining procedures, and data acquisition parameters are provided in the Supplementary Methods.

### Colony forming unit assay

Colony-forming unit (CFU) assays were performed to assess the effects of candidate compounds on progenitor activity. OCI-AML-8227 and cord blood cells were treated with compounds at concentrations selected based on prior dose-response analyses and subsequently plated in MethoCult™ H4435 Enriched medium (STEMCELL Technologies). Colonies were scored after 12 days according to standard morphological criteria. More information in Supplementary Methods.

### *Ex vivo* treatment with candidate compounds

Mouse experiments were performed under protocols approved by McGill University and affiliated research institutes. Female NSG-S mice (8–12 weeks old) were randomly assigned to treatment groups and investigators were blinded during conduct and analysis of experiments. OCI-AML- 8227 cells were treated with 500 nM BIO-acetoxime, SJB2-043, UMxxxxx03, or DMSO control for seven days prior to flow cytometric analysis or transplantation. For xenotransplantation assays, recipient mice were irradiated (2.1 Gy) 24 h prior to intrafemoral injection of treated OCI-AML- 8227 cells (1 000 CD34⁺CD38⁻ cells equivalent per DMSO control; *n* = 5 mice per condition). Twelve weeks after transplantation, bone marrow and spleen were harvested and human engraftment was assessed by flow cytometry.

For more Materials and Methods, please see: Supplemental_Material_Methods.pdf

## Results

### Low seeding density enables CD34^+^ LSC maintenance for high-throughput screening in OCI-AML-8227

To enable HTS for anti-LSC compounds using readouts such as the CellTiter Glo® Luminescence viability assay, a large number of phenotypically stable and relatively pure LSCs are required. Recent studies have identified human primary AML samples that preserve a functional LSC hierarchy *in vitro*, enabling direct investigation of LSCs (19–24). Among these, OCI-AML-8227 was selected because it maintains a stable, rare LSC-enriched CD34^+^ population with stemness and leukemia-initiating capacity, grows without stromal support and has been validated in long-term xenotransplantation assays (19–21). Whereas scaling is often trivial in established AML cell lines, the primary OCI-AML-8227 system is highly sensitive to culture conditions, with vessel format, seeding density and media composition significantly impacting CD34^+^ LSC maintenance, necessitating systematic optimization. We estimated that ∼60 million CD34⁺ LSC-enriched cells would be required to screen >11 000 compounds, necessitating ∼6×10⁹ total cells as starting material for CD34^+^ cell isolation. To expand the LSC-enriched population of OCI-AML-8227, we adapted the OCI-AML-8227 conditions from the standard 24-well format to T75 flasks. No significant differences in growth rate or CD34⁺ frequency were observed one week after transfer to T75 flasks compared with cells maintained in 24-well plates (unpaired t-test, *p* = 0.40 and *p* = 0.70). Furthermore, growth rates and CD34^+^ frequencies were maintained during an additional week of cultured in T75 flasks (unpaired t-test, *p* > 0.05), confirming scalability using larger vessels without compromising LSC content (Fig. S1A-C).

A key challenge in HTS is the spontaneous differentiation of stem cells, which can obscure LSC assays. To preserve the CD34⁺ LSC state during the six-day screening period and reduce the number of cells required, we evaluated the impact of cell density on CD34^+^ maintenance by testing AML differentiation inhibitors SR1 and UM729 (25, 26). We seeded CD34⁺ enriched cells at varying densities (100–5 000 cells/well) in 384-well plates with or without SR1 and UM729. Across all tested densities, SR1 and UM729 significantly maintained the CD34⁺ frequency and reduced differentiation markers, including CD14⁺ and CD11b⁺ blasts (Fig. S1D-H). Notably, at the lowest density tested (250 cells/well), SR1+UM729 maintained the CD34⁺ population the most effectively, nearly 15-fold over control (*p* < 0.001), suggesting that lower cell density can be used for HTS without compromising LSC maintenance (Fig. S1E).

Altogether, these optimized culture and drug assay conditions, characterized by >90% CD34⁺ purity, ∼70% recovery, maintenance over six days, and compatibility with a 384-well format at low seeding density, enabled a robust HTS directly on an LSC-enriched population from a primary AML sample. These optimized parameters formed the foundation of our large-scale compound screen.

### Primary screen identifies 61 anti-LSC inhibitors with minimal toxicity to normal hematopoietic stem and progenitor cells

To identify novel compounds that effectively target LSCs, we performed a high-throughput chemical screen of 11 142 compounds on CD34⁺ LSC-enriched OCI-AML-8227 cells (Supp. Fig. 2A and Table S1), using healthy human CD34⁺ cord blood (CB) hematopoietic stem and progenitor cells (HSPCs) as controls. The compound library included 1 960 non-FDA/FDA-approved agents with known targets (APExBIO; Table S2-S3), 4 410 investigational compounds without known targets (Table S2-S4), and 4 772 structurally diverse molecules from the Institute for Research in Immunology and Cancer (IRIC) of the Université de Montréal medicinal chemistry platform (Table S2).

Following normalization to DMSO controls, 667 compounds showed greater than 70% inhibition of CD34⁺ LSC-enriched OCI-AML-8227 cells (Fig. 1A). Of those, we selected 61 candidate hits based upon the additional criteria of toxicity to CB CD34⁺ cells (≤30%) and with the largest difference in response between OCI-AML-8227 and CB (Fig. 1B and Table S5). Among the 61 inhibitors, 22 (36%) had known targets and mechanisms, mostly from the APExBIO library, while the remaining 39 (64%) had unknown mechanisms of action (Fig. S2B and Table S5). These included known classes such as GSK3α/β inhibitors, apoptosis regulators, and HDAC inhibitors. Notably, Venetoclax, a BCL2 inhibitor with FDA approval and known anti-LSC activity, was identified among the hits, providing internal validation of our screening approach (7).

**Figure 1:**
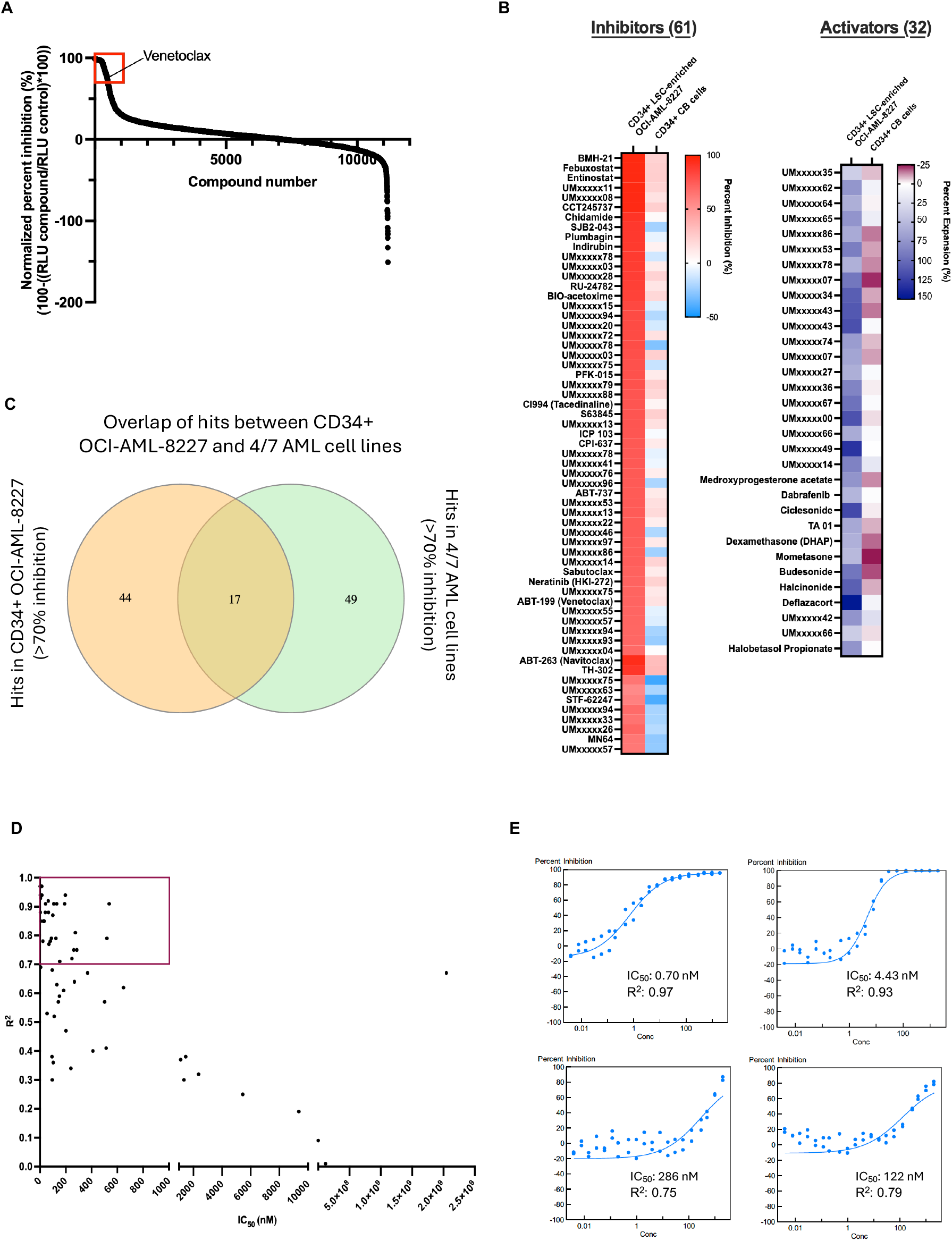
High-throughput chemical screen of 11 142 compounds and secondary validation on the CD34^+^ LSC-enriched fraction of OCI-AML-8227 identifies 33 selective modulators of LSC activity. (**A**) Waterfall plot of normalized relative luminescence units (RLU). Hits were defined as >70% inhibition in CD34^+^ LSC-enriched OCI-AML-8227 cells and <30% in CD34^+^ CB and highlighted with a red box. Venetoclax is shown. *N* = 1 biological replicate. (**B**) Activity profiles of 61 inhibitors and 32 activators in CD34^+^ LSC-enriched OCI-AML-8227 versus CD34^+^ CB cells. Inhibitors indicated by percent inhibition and activators by percent expansion of cells after treatment. (**C**) Venn diagram showing overlap between the 61 hits of CD34^+^ LSC-enriched OCI-AML-8227 and the hits from 4/7 AML cell lines (criteria: >70% inhibition in ≥50% of cell lines and <30% inhibition in two CBs). (**D**) Correlation between R^2^ values and IC_50_ (nM) for 61 inhibitors tested in secondary dose response in CD34^+^ LSC-enriched OCI-AML-8227 cells. *N* = 1 biological replicate. Compounds meeting R^2^ > 0.70 and IC_50_ < 1000 nM (red box) were retained. (**E**) Representative dose-response curves. Examples include two potent inhibitors with strong fits (top) and two moderate inhibitors (bottom).

In contrast, 32 compounds induced greater than 50% expansion of the OCI-AML-8227 cells, suggesting the potential differentiation of LSCs (Fig. 1B and Table S5). These included 22 compounds from the IRIC library and 10 from commercial libraries. Eight were steroids, mometasone, budesonide, halcinonide, and dexamethasone, previously shown to deplete LSC-enriched populations and expand AML blasts in our earlier work (19), as well as medroxyprogesterone acetate, ciclesonide, halobetasol propionate and deflazacort (Table S5). Collectively, the primary screen uncovered 61 selective anti-LSC compounds and several agents with potential LSC-differentiating activity for further investigation.

### OCI-AML-8227 uncovers LSC-specific hits and recapitulates patient sample drug responses

To assess the ability of our screen to identify compounds that would be missed in a conventional cell line screen or on bulk AML cells, we first compared the results from the seven cell lines in this screen with those from the CD34⁺ LSC-enriched OCI-AML-8227 cells (Table S1- S6). Filtering for compounds with >70% inhibition in at least 50% of cell lines and with minimal toxicity (<30% in two CBs tested) resulted in 66 cell line hits (Fig. 1C). Of our 61 anti-LSC candidates, 44 compounds were specifically anti-LSC and only 17 overlapped with the cell line hits (Fig. 1C). In addition, performing unsupervised clustering of the hits in patient samples, models and cell lines demonstrated that OCI-AML-8227 clustered with patient samples, demonstrating the utility and functionality of this model (Fig. S2C). These findings demonstrate the utility of screening directly on purified LSC populations to uncover both known and novel compounds, offering new therapeutic avenues beyond those identified through conventional bulk cell or cell line screening approaches.

### Dose response analysis refines primary hits to 33 compounds with low IC_50_ against LSC-enriched fraction

The primary screen yielded a hit ratio of 0.83%, identifying 93 initial hits, including compounds that induced expansion (Table S5). To refine this list and eliminate compounds without a dose dependent response and false positives, we performed a 20-point dose–response assay of all 93 candidates on the CD34⁺ LSC-enriched OCI-AML-8227 cells and used the CellTiter Glo® Luminescence assay to readout viability (Fig. S2D and Table S7). From this, 50 of the compounds had an R² > 0.70 and IC₅₀ < 1 000 nM and were defined as potent anti-LSC hits (Fig. 1D-E and Table S7). Thirty-three compounds demonstrated inhibitory effects (positive percent inhibition), while 17 exhibited increased total cell count (negative percent inhibition), suggesting they may promote differentiation and expansion of CD34^-^ cells (Table S7).

Of the 17 compounds that induced differentiation of CD34⁺ cells, five were structurally characterized glucocorticoids, including dexamethasone previously identified in our earlier screen (19). The additional glucocorticoids included medroxyprogesterone acetate, ciclesonide, deflazacort, and halobetasol propionate, all of which exhibited high expansion activity on CD34⁺ OCI-AML-8227 cells (Table S7). These results define a refined set of potent anti-LSC inhibitors and differentiation-inducing agents, from which we prioritized the inhibitor subset for subsequent mechanistic and functional characterization.

### Validation by flow cytometry confirms specificity of LSC targeting in AML

At screen initiation, OCI-AML-8227 cultures were ∼90% CD34⁺, but over six days of treatment the cells proliferate and partially differentiate into a mixture of CD34⁺CD38⁻ LSC-enriched cells, CD34⁺CD38⁺ leukemic progenitor cells (LPCs), and CD34⁻ differentiated blasts (∼50%) (Fig. S1D). Given that CellTiter-Glo readouts from the primary and secondary screens reflect total viability without distinguishing between subpopulations, we next assessed whether the compounds selectively target LSCs versus other AML fractions. To capture the effects of treatment across these subtypes, we performed flow cytometry on all 33 dose-validated inhibitors after six days (Fig. 2A-C). Filtering for potent agents with LC₅₀ < 500 nM in AML CD34^+^ cells reduced the panel from 33 to 24 potent candidates (Fig. 2D and Table S8). Notably, several compounds exhibited comparable LC_50_ values in CD34^+^ and CD34^-^ populations, others displayed markedly greater activity against the CD34^+^ LSC-enriched fraction, indicating differential selectivity across AML cell states (Fig. 2D and Table S8). Venetoclax again emerged as a potent hit, providing further internal validation of the screening strategy (Fig. 2D). Mechanistic annotation of the 24 compounds identified several classes including anti-apoptotic inhibitors, epigenetic inhibitors, ubiquitin protease inhibitors, among others, and compounds with unknown targets, thereby revealing multiple candidate pathways for selective LSC targeting (Fig. 2E).

**Figure 2:**
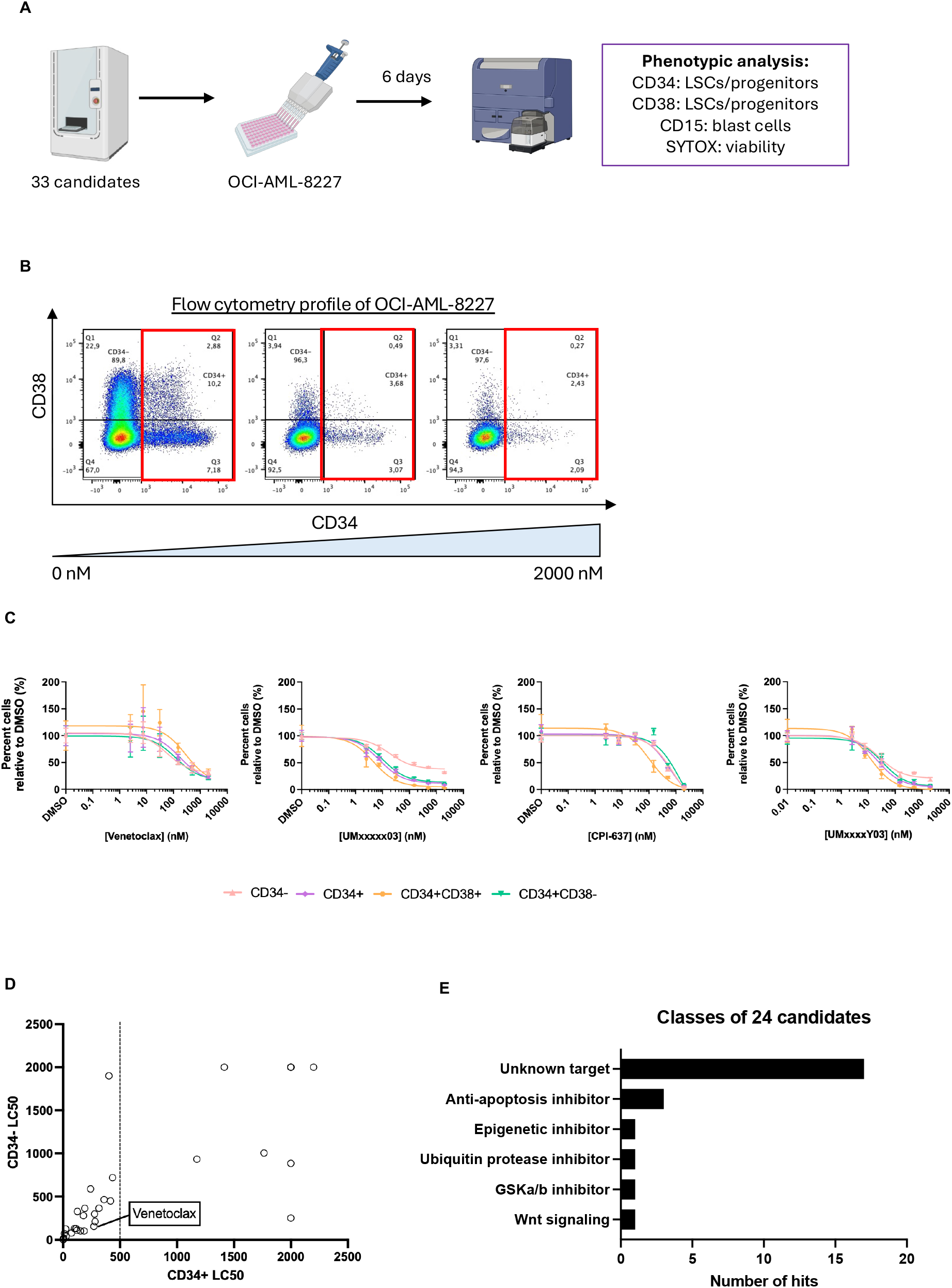
Tertiary flow cytometry validation refines candidates and confirms AML LSC specificity. **(A)** Schematic of tertiary dose response assay: 33 inhibitors were tested in OCI-AML- 8227 to assess activity across leukemic subpopulations by flow cytometry. *N* = 1 biological replicate. **(B)** Representative flow cytometry profiles of OCI-AML-8227 with increasing drug concentration, illustrating CD34^+^ (red box) reduction. **(C)** Dose response curves for representative inhibitors across CD34^-^ blasts and CD34^+^ subpopulations of OCI-AML-8227. Mean ± s.d. **(D)** Plot of CD34⁺ vs CD34⁻ LC₅₀ values; compounds with LC₅₀ < 500 nM in LSCs were prioritized. Venetoclax is highlighted. **(F)** Classification of 24 refined hits: 17 unknown targets, seven annotated inhibitors.

### Candidates induce apoptosis in LSC-enriched fractions in a second LSC model

Twenty of the 24 identified compounds are not FDA approved or in preclinical trials for AML, making them compelling candidates for further evaluation. As an intermediate step to prioritize hits for downstream mechanistic analyses, we first assessed whether their activity extended beyond OCI-AML-8227 and could be maintained in a microenvironment-dependent model. We evaluated the twenty compounds in OCI-AML-20, a poor-prognosis therapy-refractory LSC model that requires OP9 stromal co-culture for CD34⁺ maintenance and harbors an inv(3)/*EVI1* lesion (22) (Fig. 3A-B). Five inhibitors produced ≥50% reduction in the CD34⁺ LSC-enriched fraction while remaining non-toxic to stromal cells (Fig. 3C and Fig. S3A). LC₅₀ values for these five inhibitors in CD34⁺ OCI-AML-20 cells were similar to, or modestly higher than, those in OCI-AML-8227 (Table S9). Although overall sensitivity was lower in OCI-AML-20, robust depletion of LSCs in this stroma-supported model enabled prioritization of a subset of candidates for subsequent apoptosis-focused analyses and confirms that several hits retain activity against aggressive AML with a protective niche, underscoring their translational potential.

**Figure 3:**
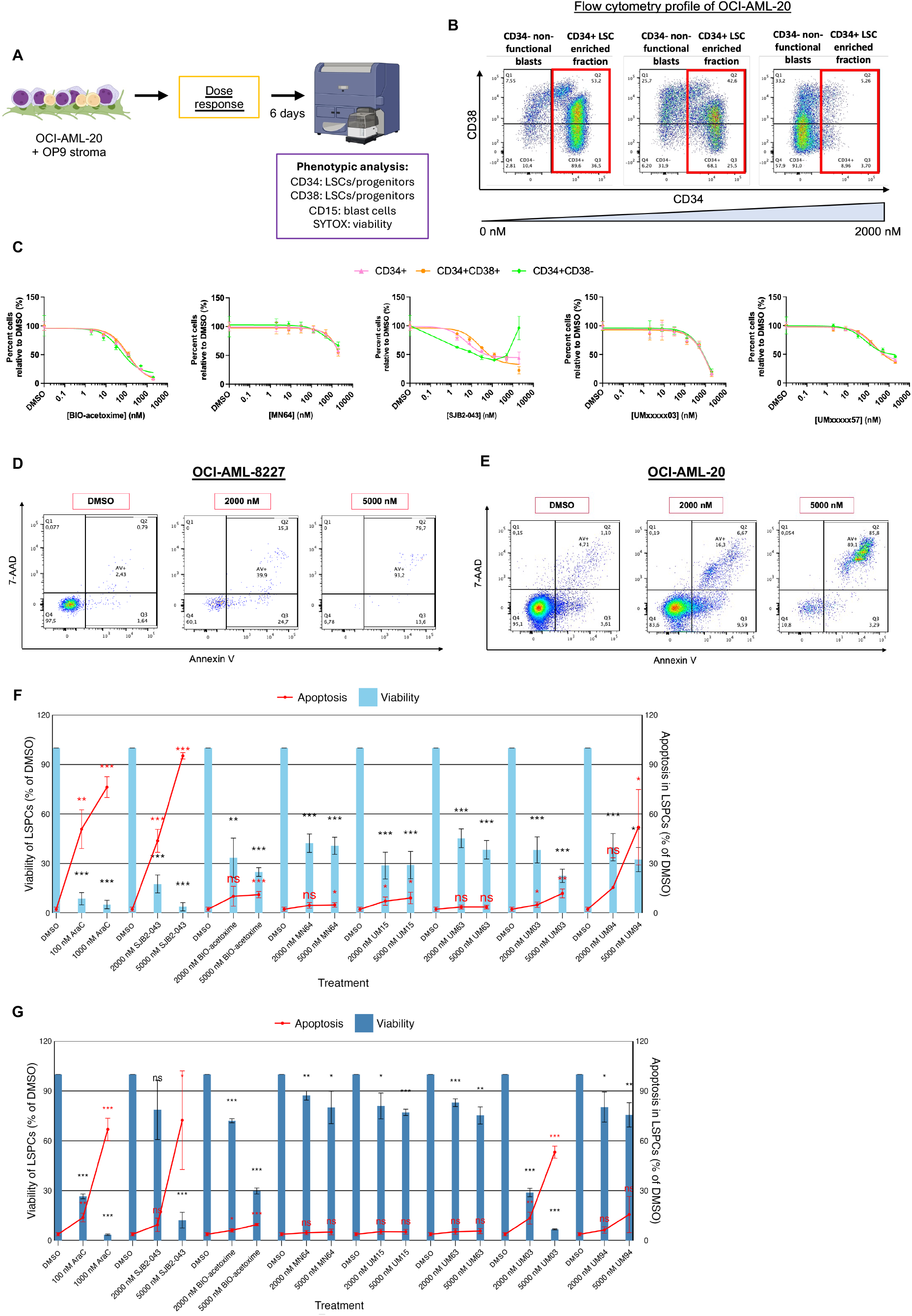
Validation of lead compounds in a second poor-prognosis LSC model (OCI-AML- 20) and assessment of apoptosis. **(A)** Schematic of dose response in OCI-AML-20 cells co-cultured with OP9 stromal cells. Cells were treated for six days and analyzed by flow cytometry. **(B)** Representative CD38 vs CD34 flow cytometry profiles of OCI-AML-20 treated with increasing doses of BIO-acetoxime. Red box highlights CD34^+^ LSC-enriched population. **(C)** Dose–response curves of OCI-AML-20 for five lead compounds, showing effects on total CD34⁺ (total LSC fraction), CD34⁺CD38⁺ (LPC-enriched), and CD34⁺CD38⁻ (LSC-enriched) fractions. Mean ± s.d. Representative one of *n* = 3 biological replicates. **(D-E)** Representative 7-ADD vs Annexin V flow cytometry plots in **(D)** OCI-AML-8227 and **(E)** OCI-AML-20 for CD34^+^ cells treated with SJB2-043. **(F-G)** Viability of CD34^+^ cells normalized to DMSO (bars) and apoptosis (lines) in CD34^+^ fractions of OCI-AML-8227 **(F)** and **(G)** OCI-AML-20 treated with lead compounds and Ara-C control for three days. Mean ± s.d.; unpaired t-test, \**p* < 0.05, \*\**p* < 0.01, \*\*\**p* < 0.001, \*\*\*\**p* < 0.0001). Apoptosis significance: >5% AV+ change and p-values. Representative is *n* = 4 biological replicates combined for both samples.

To determine whether prioritized candidates deplete LSC-enriched populations through apoptosis, we evaluated seven top hits based on their activity against CD34⁺ populations in OCI-AML-8227 and/or OCI-AML-20, and low toxicity toward stromal cells. Apoptosis was assessed in the CD34^+^ fraction using Annexin V staining, with a threshold of >5% increase compared to untreated cells and a significance threshold of *p* < 0.05 (Fig. 3D-E). BIO-acetoxime, UMxxxxx03, UMxxxxx15, UMxxxxx94, and SJB2-043 induced apoptosis in CD34^+^ cells in either OCI-AML- 8227 or OCI-AML-20 (Fig. 3F-G). Together, these results demonstrate that a subset of prioritized compounds depletes LSC-enriched populations through induction of apoptosis in adverse-risk AML.

### Primary patient LSC-enriched populations are eliminated by candidate compounds

To verify whether compounds can target LSCs in a wide range of primary AML samples and if this eradication also occurs through apoptosis, the seven candidates were tested on nine primary patient samples obtained from the Quebec Leukemia Cell Bank (BCLQ) and primary, uncultured OCI-AML-8227 patient sample (Fig. 4A). These included six intermediate risk samples and four adverse risk samples (nine samples + OCI-AML-8227) and contained multiple mutations confirmed by next generation sequencing (Fig. S4A-C and Table S10). All seven candidates were able to target the CD34^+^ and CD34^+^CD38^-^ LSC-enriched fractions in most samples with BIO-acetoxime, SJB2-043 and UMxxxxx03 inducing a response in 9/10 samples tested (Fig. 4B and Table S10). Apoptosis in CD34⁺ cells was confirmed for BIO-acetoxime, UMxxxxx03, UMxxxxx94, and SJB2-043 in most patient samples (Fig. 4C). Overall, we identified candidate compounds that eliminate LSC-enriched cells in most primary intermediate and adverse risk AML samples, partially through apoptosis.

**Figure 4:**
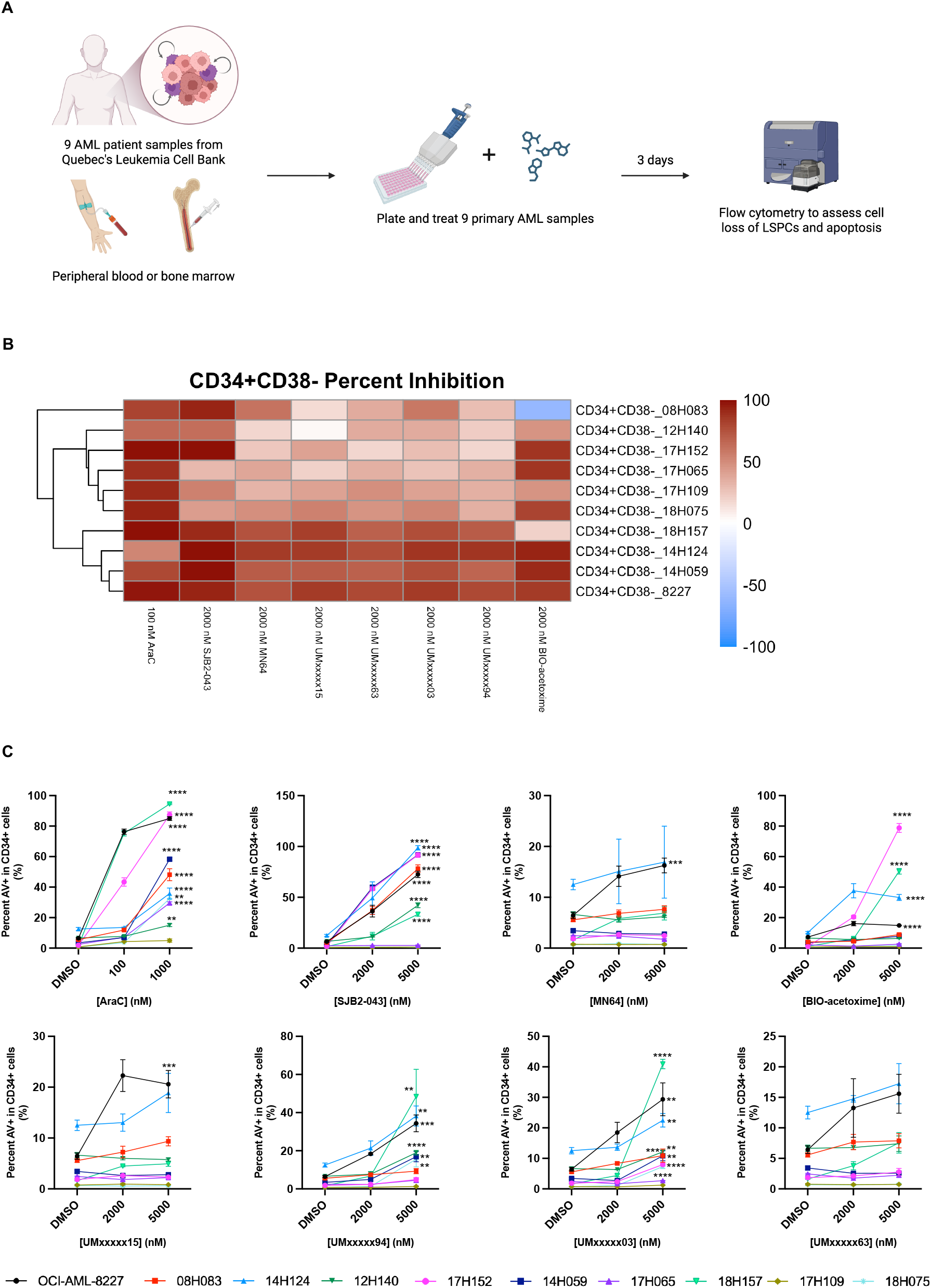
Top seven candidates eliminate CD34^+^ fractions in primary AML patient samples via apoptosis. **(A)** Schematic of treating primary patient samples (*n* = 9) from the BCLQ cohort with seven candidate compounds and cytarabine as a positive control for three days. Samples consist of peripheral blood or bone marrow. Samples were acquired by flow cytometry, and apoptosis was measured using Annexin V and 7-AAD staining. *N* = 1 biological replicate. **(B)** Heatmap showing percent inhibition of CD34^+^CD38^-^ LSC-enriched fractions of nine primary AML patient samples after three days of treatment with candidate at 2 000 nM. Higher inhibition reflects greater loss of population. Cytarabine was used as a control at 100 nM. *N* = 1 biological replicate. **(C)** Apoptosis in CD34^+^ fractions of nine primary patient samples measured by flow cytometry. Apoptosis was defined as >5% increase vs. untreated; significance was determined by unpaired t-test, \**p* < 0.05, \*\**p* < 0.01, \*\*\**p* < 0.001, \*\*\*\**p* < 0.0001). Mean ± s.d. *N* = 1 biological replicate.

### Top candidates target LPCs while sparing normal progenitors *in vitro*

Based on their consistent activity in eliminating the LSC-enriched fractions of primary samples and ability to induce apoptosis, we prioritized three candidates for further evaluation of selectivity toward normal HSPCs. Although toxicity towards healthy HSPCs was included as a filter in the initial screen (<30% inhibition), this assessment was based on bulk measurements that do not resolve specific cell populations. Therefore, to directly evaluate selectivity at the cellular level, CD34^+^ CB cells and OCI-AML-8227 were treated with three of the lead candidates: two candidates with known targets (BIO-acetoxime and SJB2-043) and one with an unknown mechanism (UMxxxxx03). Mild toxicity was observed only at higher doses (>500 nM), and LC₅₀ values for HSPCs were approximately 20- to 100-fold higher than those for the CD34⁺ LSC-enriched fraction in OCI-AML-8227 (Fig. 5A and Fig. S5A-B). These findings confirm that LSCs can be selectively targeted at doses that spare healthy HSPCs.

**Figure 5:**
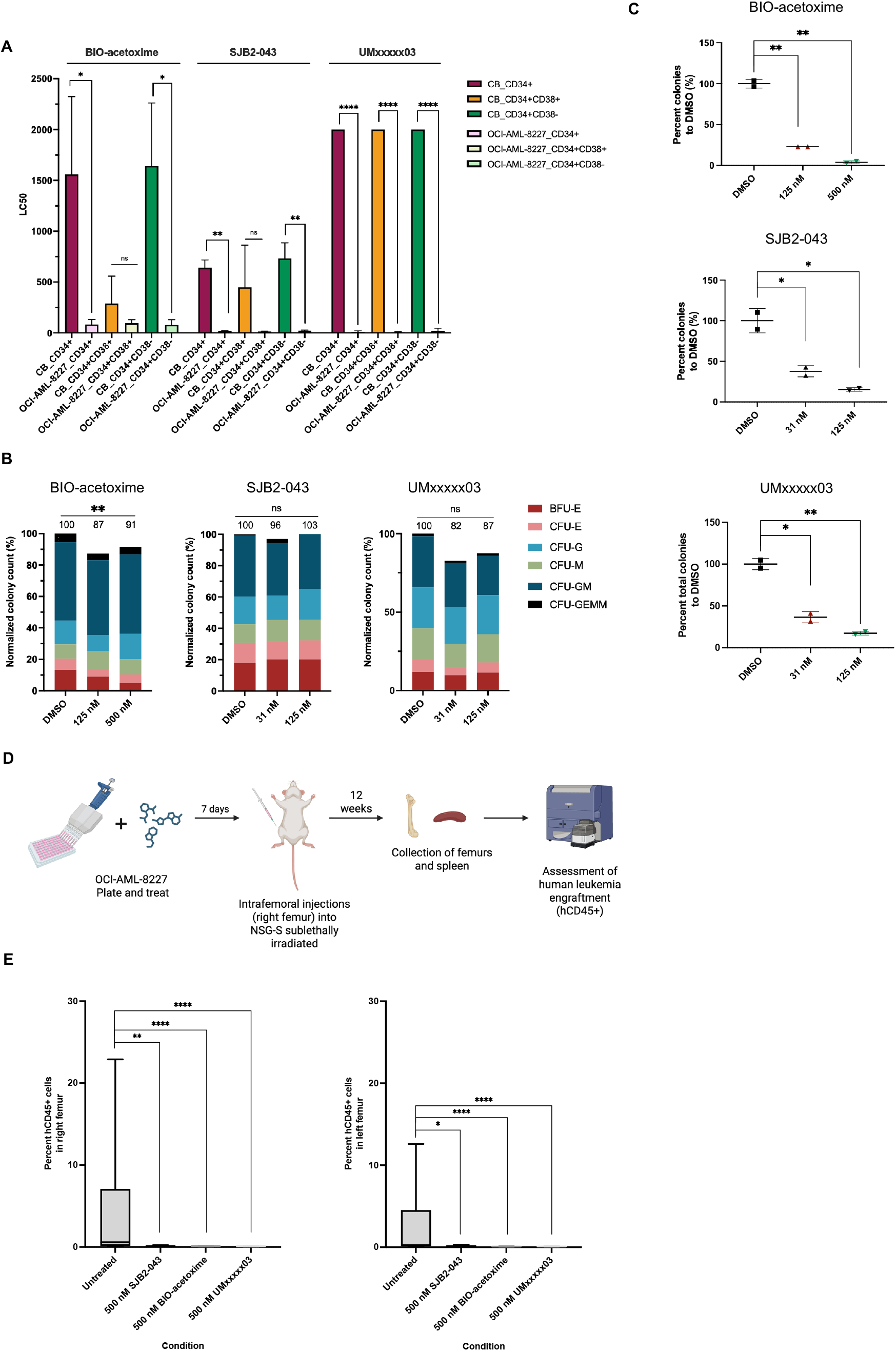
Three lead compounds reduce LSC clonogenic potential *in vitro* and leukemia initiating potential *in vivo*. **(A)** Summary of LC_50_ from three lead candidates BIO-acetoxime, SJB2-043 and UMxxxxx03 on CD34^+^ CB and OCI-AML-8227 subpopulations. *N* = 3 biological replicates. **(B)** CFU assay of healthy CD34^+^ CB HSPCs treated for six days with candidate compounds at doses above the LC₅₀ of leukemic cells. Colony number was normalized to DMSO and counted at 12 days. Representative one of *n* = 3 biological replicates. Data are mean ± s.d.; \**p* < 0.05, \*\**p* < 0.01, \*\*\**p* < 0.001, \*\*\*\**p* < 0.0001 (unpaired t-test). **(C)** CFU assay of OCI-AML- 8227 cells treated for six days with candidate compounds at their LC₅₀. Colony number was normalized to DMSO and counted at 12 days. Representative one of *n* = 3 biological replicates. Data are mean ± s.d.; \**p* < 0.05, \*\**p* < 0.01, \*\*\**p* < 0.001, \*\*\*\**p* < 0.0001 (unpaired t-test). **(D)** Experimental schematic for *in vivo* xenotransplantation assay. Bulk OCI-AML-8227 cells were treated for seven days and then 1 000 CD34⁺CD38⁻ cells were injected intrafemorally into sublethally irradiated (2.1 Gy) NSG-S mice. After 12 weeks, femurs were harvested for analysis. **(E)** Human CD45⁺ engraftment in injected (right) and contralateral (left) femurs at 12 weeks. *N* = 2 biological replicates, *n* = 10 mice per group. Paired t-test; *p* < 0.05, \*\**p* < 0.01, \*\*\**p* < 0.001, \*\*\*\**p* < 0.0001.

To determine whether prioritized candidates target functional LPCs while sparing normal HSPCs, we performed colony forming unit (CFU) assays on OCI-AML-8227 and CB cells following six days of treatment (Fig. 5B-C and Fig. S5C-D). All three candidates significantly reduced AML progenitor colony formation, whereas CB colony-forming capacity (CFU-E, BFU-E, CFU-GM, CFU-G, CFU-M, CFU-GEMM) remained largely unaffected across conditions (Fig. 5B-C). These findings define a therapeutic window in which LPCs are eliminated while normal hematopoietic progenitors are preserved.

### Candidates eliminate LSC leukemia initiating potential

To confirm that candidates target *bona fide* functional LSCs, we performed long-term LSC xenotransplantation assays with the top three candidates (BIO-acetoxime, SJB2-043 and UMxxxxx03) based on their potency *in vitro* and ability to achieve robust LPC reduction at lower concentrations (<500 nM). Leukemic cells were treated for seven days and injected intrafemorally into the right femurs of NSG-S mice (Fig. 5D and Fig. S5E-F). After 12 weeks, human engraftment (hCD45^+^) in the injected and contralateral femurs was almost eliminated following treatment with each compound compared to untreated controls (Fig. 5E), confirming that these candidates effectively target functional LSCs.

### Single-cell profiling reveals convergent stress responses with distinct functional outcomes in LSCs

To define the transcriptional programs underlying LSC depletion and candidate mechanism of action, we performed single-cell RNA sequencing (scRNA-seq) on OCI-AML-8227 cells treated with the three top candidates for 24 hours: SJB2-043, BIO-acetoxime, and UMxxxxx03 (Fig. 6A and Fig. S6A-C). For each compound, transcriptional profiling revealed loss of stemness, with a marked reduction in the LSC population observed by both UMAP and flow cytometry (Fig. 6A-B and Fig. S6D-F), and decreased expression of stemness associated genes (Fig. 6C and Table S11-S13). Additionally, the multi-lineage GMP cluster had significant negative enrichment of stemness signatures, further supporting disruption of stem-like transcriptional states across populations (Fig. S6G-I).

**Figure 6:**
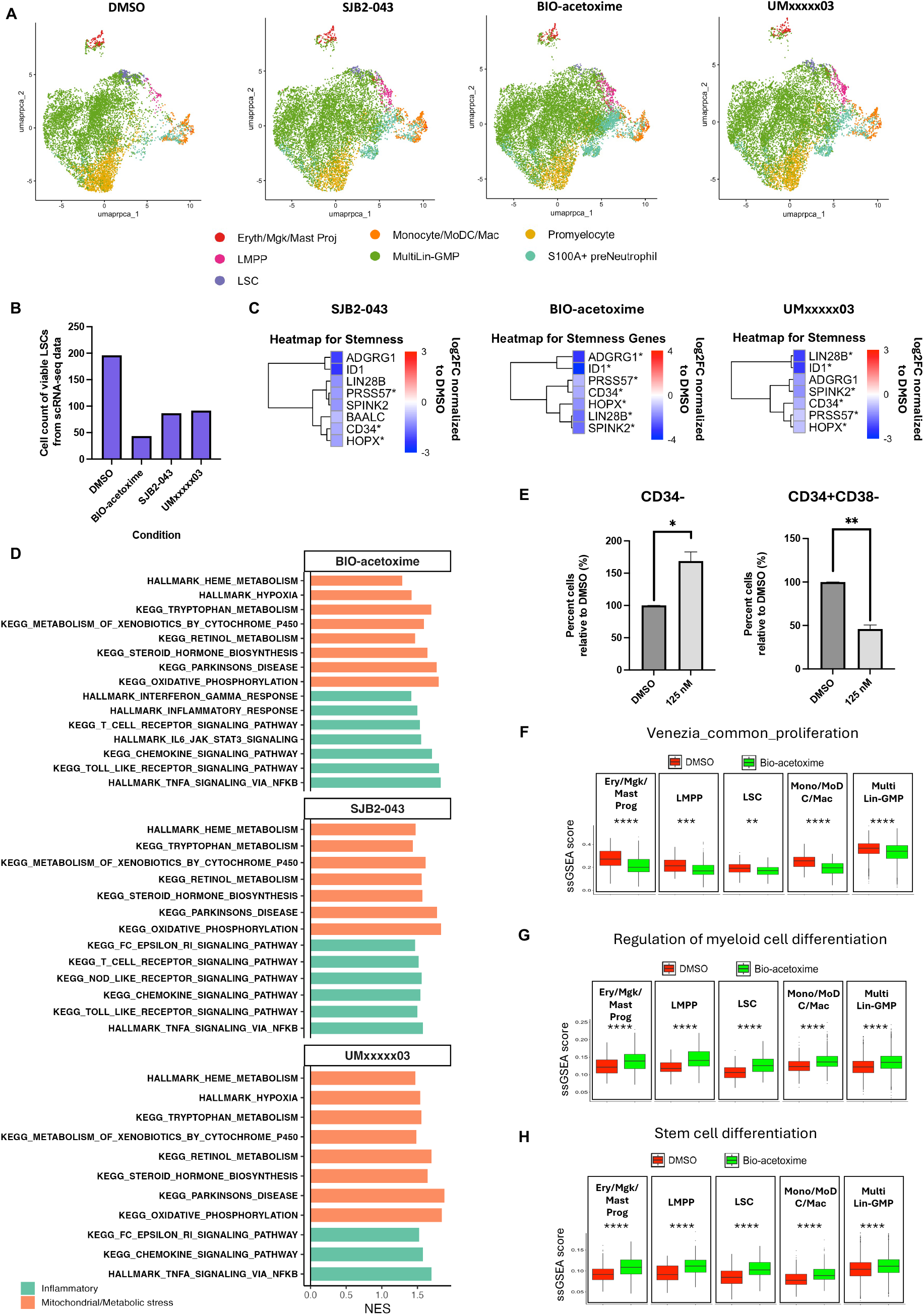
Single-cell transcriptional profiling reveals reduction in stemness and convergence on metabolic and inflammatory stress responses in LSCs after treatment with lead compounds. **(A)** UMAP plots of CD34⁺-enriched OCI-AML-8227 cells after 24 h treatment with 250 nM SJB2-043, 500 nM BIO-acetoxime, or 1 000 nM UMxxxxx03. Clusters were annotated using the ANNCAST classifier v1.2 into seven populations: LSC, lympho-myeloid progenitors (LMPP), multi lineage granulocyte-monocyte progenitors (MultiLin-GMP), monocyte/macrophage/DC lineage (Monocyte/MoDC/Mac), promyelocyte, S100A+ preNeutrophil, erythroid/megakaryocyte/mast progenitors (Ery/MK/Mast Proj). *N* = 1 biological replicate for scRNA-seq data. **(B)** Quantification of viable LSC cell count from LSC cluster scRNA-seq data is shown for each condition. **(C)** Heatmaps of stemness-associated genes in LSC clusters after treatment with candidate compounds. **(D)** GSEA of LSC clusters with metabolic stress (orange) and inflammatory signaling (green) pathways. Signatures with *p* < 0.05 and FDR < 0.25 are shown; NES = normalized enrichment score. **(E)** Normalized viable cell counts to DMSO of OCI-AML-8227 CD34^-^ (blasts) and CD34^+^ (LSC-enriched fraction) after six-day treatment with 125 nM BIO-acetoxime. Data are mean ± s.d. (*n* = 3 biological replicates); \**p* < 0.05, \*\**p* < 0.01, \*\*\**p* < 0.001, \*\*\*\**p* < 0.0001 (unpaired t-test). **(F–H)** ssGSEA of the LSC cluster of BIO-acetoxime–treated cells (24 hr) with HSPC proliferative and stem/myeloid differentiation signatures. Statistical significance of ssGSEA enrichment scores was calculated using a two-sided Wilcoxon test. Genes with an asterisk are significant and have *p* < 0.05.

Despite targeting distinct molecular pathways, enrichment analysis of the LSC cluster revealed that all three compounds converge on core transcriptional programs associated with metabolic stress and inflammatory signaling (Fig. 6D and Table S14-S16). These convergent stress-response programs point to metabolic and redox imbalance as a shared vulnerability in LSCs. Despite the shared signatures, response to the compounds are functionally resolved in distinct ways: SJB2-043 and UMxxxxx03 promote apoptotic priming (Fig. 3F, G), whereas BIO-acetoxime predominantly drives differentiation (Fig. 6E-H). Together, these findings support a model in which common stress-induced transcriptional programs are differentially leveraged to drive distinct outcomes in LSCs. Given its potent and selective anti-LSC activity, we prioritized SJB2-043 for further mechanistic characterization.

### SJB2-043 eradicates LSCs through USP1 inhibition, leading to oxidative stress-induced apoptosis

SJB2-043 is a known USP1 inhibitor with reported anti-leukemia activity, although its effect on LSCs has not been explored (27). Consistent with target engagement, treatment with SJB2-043 significantly reduced USP1 protein levels in CD34^+^ LSC-enriched OCI-AML-8227 cells and was accompanied by loss of *ID1*, a key regulator of LSC self-renewal identified by scRNA-seq (Fig. 7A-B). SJB2-043 treatment also decreased expression of *FANCD2*, a critical effector of the Fanconi anemia (FA) DNA repair pathway and established USP1 substrate (Fig. 7B) (28).

**Figure 7:**
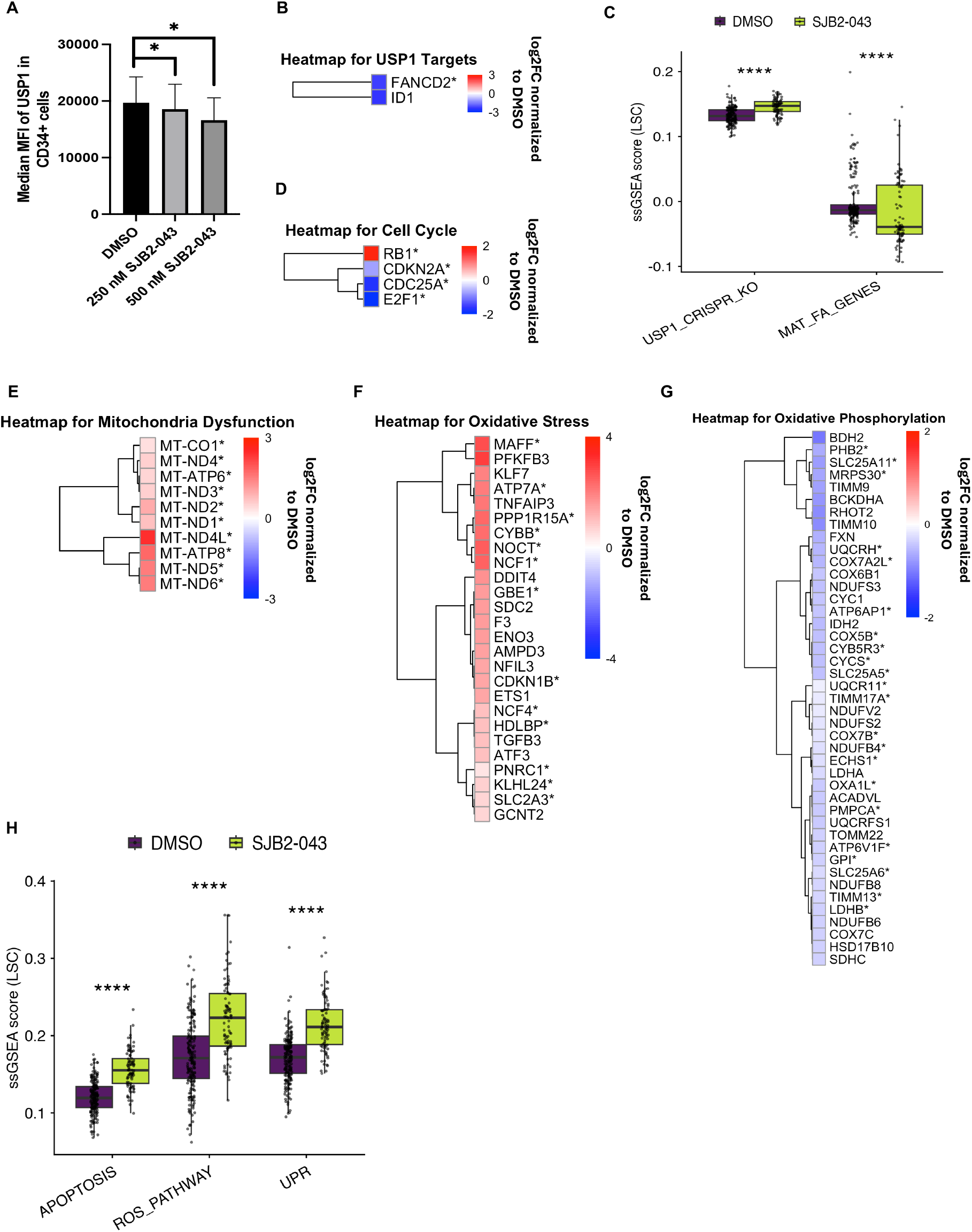
Mechanism of action of SJB2-043 for LSC eradication. **(A)** Intracellular flow cytometry showing USP1 protein levels in CD34⁺ OCI-AML-8227 cells after 24 h treatment with SJB2-043. MFI: Median fluorescence intensity. Data are mean ± s.d. (*n*=4 biological replicates); one-way ANOVA with multiple comparisons, \**p* < 0.05, \*\**p* < 0.01, \*\*\**p* < 0.001, \*\*\*\**p* < 0.0001. **(B)** Heatmap of FC normalized to DMSO for USP1 targets in the LSC cluster following SJB2-043 treatment (250 nM). **(C)** ssGSEA of LSC cluster cells with the USP1 knockout transcriptional signature and Fanconi anemia DNA repair pathway. **(D)** Heatmap of FC normalized to DMSO for cell cycle regulators repressed by SJB2-043 in the LSC cluster. **(E-G)** Heatmaps of FC normalized to DMSO of leading-edge genes of mitochondrial dysfunction **(E)**, oxidative stress **(F)** and oxidative phosphorylation **(G)** in the LSC cluster after SJB2-043 treatment (250 nM). **(H)** ssGSEA of stress programs in LSC cluster treated with 250 nM SJB2-043 (24 hr). Statistical significance of ssGSEA enrichment scores was calculated using a two-sided Wilcoxon test; genes with an asterisk are significant and have *p* < 0.05. All scRNA-seq data is *n* = 1 biological replicate.

Given USP1’s role in regulating FANCD2 and FANCI within the FA pathway, we assessed the expression of a USP1 CRISPR knockout (KO) transcriptional signature and an FA pathway gene set (Fig. 7C). SJB2-043 treatment resulted in significant positive enrichment of the USP1 KO signature, consistent with transcriptional mimicry of USP1 loss, and significant negative enrichment of the FA pathway signature, indicating suppression of Fanconi anemia-mediated DNA repair (Fig. 7C). Given the observed impairment of DNA repair and the role of RB1 in stress-induced cell cycle regulation (29), we next examined key cell cycle regulators (Table S11). Multiple G1/S regulators, including *CDC25A*, *CDKN2A*, and *E2F1*, were repressed, suggesting attenuation of cell cycle progression and reduced proliferative capacity (Fig. 7D).

Consistent with the role of USP1 in genome stability and cell cycle control, we then examined mitochondrial and translational stress responses downstream of DNA repair disruption. Single sample gene set enrichment analysis (ssGSEA) and GSEA indicated that SJB2-043 treatment induced pronounced mitochondrial stress, with dysregulation of mitochondrial genes, oxidative phosphorylation genes, and redox balance components, indicative of ETC imbalance and mitochondrial dysfunction (Fig. 7E-G). Translational stress was further supported by upregulation of *EIF4G2* and *EIF4B* and downregulation of ribosomal genes (Table S11 and S14).

Treatment with SJB2-043 also induced inflammatory signaling, including upregulation of *STAT1*, *IL32*, *CXCL8*, and other inflammation-linked genes (Fig. S6A and Table S14), together with ssGSEA showing positive enrichment in hallmark unfolded protein response, ROS and apoptosis signatures (Fig. 7H). While apoptosis was confirmed functionally in cell-based assays, its transcriptional enrichment supports the conclusion that SJB2-043 triggers stress-induced apoptotic priming in LSCs (Fig. 7H).

Together, these findings support a model in which USP1 inhibition by SJB2-043 disrupts DNA repair and induces mitochondrial and oxidative stress responses that drive apoptotic priming and elimination of LSCs. These results establish USP1 as a therapeutic vulnerability in AML LSCs and provide mechanistic insight into the anti-LSC activity of SJB2-043.

## Discussion

Despite therapeutic advances, outcome is poor in AML and is driven by LSCs, a therapy-resistant subpopulation capable of self-renewal. While LSCs are an appealing therapeutic target, their rarity, purification challenges, and limited *ex vivo* expansion have hindered direct HTS. Most prior screens have focused on bulk AML cells, surrogate models, or a small number of compounds, often failing to uncover compounds with true anti-LSC activity (11–15, 19). Here, we overcame these barriers by developing a scalable HTS platform directly on a highly purified, expandable LSC-enriched population derived from OCI-AML-8227. Our screen of 11 142 structurally diverse compounds identified 20 anti-LSC hits, including known LSC-targeting drugs like venetoclax, validating the approach and enabling discovery of novel, mechanistically distinct compounds (7).

Our comparative analysis of the OCI-AML-8227 model with 34 AML samples showed that it clusters most closely with MLL-rearranged and stem-like AMLs, rather than standard cell lines (30). This suggests that screening on OCI-AML-8227 captures vulnerabilities relevant to aggressive, undifferentiated subtypes. Although some hits were shared with cell lines, nearly 72% were unique to LSCs, highlighting the utility of this screening approach for uncovering LSC-specific targets and emphasizing the distinct biology and vulnerabilities of LSCs compared to bulk.

Mechanistically, transcriptomic and functional analyses of three top LSC-depleting candidates SJB2-043, BIO-acetoxime and UMxxxxx03 converged on three broad biological themes: suppression of stemness programs, induction of metabolic and oxidative stress, and disruption of inflammatory signaling that tips cells toward apoptosis or enforced differentiation. This convergence is consistent with prior studies demonstrating that effective LSC-targeting therapies simultaneously disrupt stemness and mitochondrial metabolism while promoting oxidative and inflammatory stress. For example, Venetoclax, a BCL-2 inhibitor with clinical efficacy in AML, eradicates LSCs in part by impairing oxidative phosphorylation and mitochondrial fitness, thereby targeting a key metabolic dependency of LSCs (7, 8). Similarly, parthenolide can induce reactive oxidative species through inhibition of NF-κB signaling, leading to apoptosis in LSCs through combined oxidative and inflammatory stress (31, 32). Finally, epigenetic regulators such as BRD4 have been linked to maintenance of stemness and inflammatory transcriptional programs, and emerging evidence suggests that BRD4 inhibition can intersect with AHR-mediated pathways to suppress leukemic self-renewal and promote differentiation (18, 33). Collectively, these data suggest that these three axes represent core dependencies in LSC biology that can be exploited through diverse therapeutic strategies, such as those identified in this study.

SJB2-043, a known USP1 inhibitor, exhibited the strongest pro-apoptotic activity and selectively eradicated CD34⁺CD38⁻ cells across models. USP1 has been implicated in genome stability and maintaining stemness through *ID1* regulation, but its role in LSCs has not been defined (34–38). In ovarian cancer, USP1 mediates platinum resistance and dissemination via stabilization of the EMT regulator Snail, and in hepatocellular carcinoma it promotes progression through the Hippo/TAZ axis (39, 40). Our findings provide the first evidence that disrupting USP1 in AML eradicates functional LSCs by destabilizing genome integrity, redox balance and metabolic homeostasis. Given USP1’s regulation of FANCD2/FANCI in the FA pathway, these findings, together with our observation of FA pathway modulation following SJB2-043 treatment in AML LSCs supports further investigation of USP1 inhibition as a targeted strategy in this disease and central regulator of LSC fitness.

SJB2-043 also presents an intriguing mechanism in LSCs. While a previous work by Mistry *et al.* 2013 showed that SJB2-043 degrades USP1 and ID1 in bulk AML, our findings indicate that it targets LSCs specifically and operates through proteasomal degradation rather than classical enzymatic inhibition (27). This aligns with recent findings by Rennie *et al.*, 2024, who showed that SJB2-043 does not engage USP1’s catalytic pocket in the same manner as other deubiquitinate inhibitors (41). Our data support a non-canonical mode of action in LSCs, possibly involving broader proteostasis disruption and redox collapse, which remains to be investigated.

In summary, we developed and applied the first large-scale HTS directly on primary LSCs, leading to the identification of multiple novel compounds that selectively eliminate LSCs through diverse mechanisms. This approach uncovered SJB2-043, a USP1 inhibitor with a previously unrecognized ability to target LSCs via DNA repair disruption and oxidative stress. Additional hits like BIO-acetoxime and UMxxxxx03 engage inflammatory and metabolic pathways to induce LSC differentiation or collapse biosynthetic programs. By integrating transcriptomics, protein analysis, and functional assays, we demonstrate the value of LSC-directed screening and identified new therapeutic avenues that may ultimately improve outcomes for patients with AML.

## Supporting information

Supplemental Table Titles and Legends

## Acknowledgments

This work was in part supported by grants from Oncopole (EMC2 grant through the Fonds de recherche du Québec– Santé (FRQS)) (BTW, FB, SC, KE), the Canada Research Chair program (KE), as well as Canadian Institutes of Health Research (CIHR) and FRQS fellowships (IAI). Supported by a Blood Cancer Research Jump Start Grant from The Leukemia & Lymphoma Society of Canada / Societe de Leucemie & Lymphome Du Canada (KE). The Banque de Cellules Leucémiques du Québec is supported by grants from the Cancer Research Network of the Fonds de Recherche du Québec – Santé.

## Author contributions

Conceptualization: IAI, KE. Methodology: IAI, SSTH, SC, FB, BTW, KE. Investigation: IAI, SSTH, HD, MS, BJ, PAT, MB, ALN. Formal analysis: IAI, SSTH. Visualization: IAI, KE. Funding acquisition: IAI, SC, FB, BTW, KE. Supervision: KE. Writing – original draft: IAI, KE. Writing – review & editing: IAI, SSTH, HD, BJ, PAT, MB, ALN, JH, FB, KE.

## Data and materials availability

Transcriptomic data is prepared and will be uploaded to a public database before publishing.

## References

1. DiNardo CD, Cortes JE. Mutations in AML: prognostic and therapeutic implications. Hematology American Society of Hematology Education Program. 2016;2016(1):348–55.

2. Short NJ, Konopleva M, Kadia TM, Borthakur G, Ravandi F, DiNardo CD, et al. Advances in the Treatment of Acute Myeloid Leukemia: New Drugs and New Challenges. Cancer discovery. 2020;10(4):506–25.

3. Terwijn M, Zeijlemaker W, Kelder A, Rutten AP, Snel AN, Scholten WJ, et al. Leukemic stem cell frequency: a strong biomarker for clinical outcome in acute myeloid leukemia. PLoS One. 2014;9(9):e107587.

4. Ho TC, LaMere M, Stevens BM, Ashton JM, Myers JR, O’Dwyer KM, et al. Evolution of acute myelogenous leukemia stem cell properties after treatment and progression. Blood. 2016;128(13):1671–8.

5. Guzman ML, Neering SJ, Upchurch D, Grimes B, Howard DS, Rizzieri DA, et al. Nuclear factor-kappaB is constitutively activated in primitive human acute myelogenous leukemia cells. Blood. 2001;98(8):2301–7.

6. Lagadinou ED, Sach A, Callahan K, Rossi RM, Neering SJ, Minhajuddin M, et al. BCL- 2 inhibition targets oxidative phosphorylation and selectively eradicates quiescent human leukemia stem cells. Cell stem cell. 2013;12(3):329–41.

7. Pollyea DA, Stevens BM, Jones CL, Winters A, Pei S, Minhajuddin M, et al. Venetoclax with azacitidine disrupts energy metabolism and targets leukemia stem cells in patients with acute myeloid leukemia. Nature medicine. 2018;24(12):1859–66.

8. Jones CL, Stevens BM, D’Alessandro A, Reisz JA, Culp-Hill R, Nemkov T, et al. Inhibition of Amino Acid Metabolism Selectively Targets Human Leukemia Stem Cells. Cancer cell. 2019;35(2):333–5.

9. DiNardo CD, Jonas BA, Pullarkat V, Thirman MJ, Garcia JS, Wei AH, et al. Azacitidine and Venetoclax in Previously Untreated Acute Myeloid Leukemia. New England Journal of Medicine. 2020;383(7):617–29.

10. DiNardo CD, Pratz K, Pullarkat V, Jonas BA, Arellano M, Becker PS, et al. Venetoclax combined with decitabine or azacitidine in treatment-naive, elderly patients with acute myeloid leukemia. Blood. 2019;133(1):7–17.

11. Hassane DC, Guzman ML, Corbett C, Li X, Abboud R, Young F, et al. Discovery of agents that eradicate leukemia stem cells using an in silico screen of public gene expression data. Blood. 2008;111(12):5654–62.

12. Sachlos E, Risueño RM, Laronde S, Shapovalova Z, Lee JH, Russell J, et al. Identification of drugs including a dopamine receptor antagonist that selectively target cancer stem cells. Cell. 2012;149(6):1284–97.

13. Sykes DB, Kfoury YS, Mercier FE, Wawer MJ, Law JM, Haynes MK, et al. Inhibition of Dihydroorotate Dehydrogenase Overcomes Differentiation Blockade in Acute Myeloid Leukemia. Cell. 2016;167(1):171–86.e15.

14. Subedi A, Liu Q, Ayyathan DM, Sharon D, Cathelin S, Hosseini M, et al. Nicotinamide phosphoribosyltransferase inhibitors selectively induce apoptosis of AML stem cells by disrupting lipid homeostasis. Cell stem cell. 2021.

15. McDermott SP, Eppert K, Notta F, Isaac M, Datti A, Al-awar R, et al. A small molecule screening strategy with validation on human leukemia stem cells uncovers the therapeutic efficacy of kinetin riboside. Blood. 2012;119(5):1200–7.

16. Hartwell KA, Miller PG, Mukherjee S, Kahn AR, Stewart AL, Logan DJ, et al. Niche- based screening identifies small-molecule inhibitors of leukemia stem cells. Nat Chem Biol. 2013;9(12):840–8.

17. Dal Bello R, Pasanisi J, Joudinaud R, Duchmann M, Pardieu B, Ayaka P, et al. A multiparametric niche-like drug screening platform in acute myeloid leukemia. Blood cancer journal. 2022;12(6):95.

18. Zuber J, Shi J, Wang E, Rappaport AR, Herrmann H, Sison EA, et al. RNAi screen identifies Brd4 as a therapeutic target in acute myeloid leukaemia. Nature. 2011;478(7370):524–8.

19. Laverdiere I, Boileau M, Neumann AL, Frison H, Mitchell A, Ng SWK, et al. Leukemic stem cell signatures identify novel therapeutics targeting acute myeloid leukemia. Blood cancer journal. 2018;8(6):52.

20. Eppert K, Takenaka K, Lechman ER, Waldron L, Nilsson B, van Galen P, et al. Stem cell gene expression programs influence clinical outcome in human leukemia. Nature medicine. 2011;17(9):1086–93.

21. Lechman ER, Gentner B, Ng SW, Schoof EM, van Galen P, Kennedy JA, et al. miR-126 Regulates Distinct Self-Renewal Outcomes in Normal and Malignant Hematopoietic Stem Cells. Cancer cell. 2016;29(2):214–28.

22. Luciani GM, Xie L, Dilworth D, Tierens A, Moskovitz Y, Murison A, et al. Characterization of inv(3) cell line OCI-AML-20 with stroma-dependent CD34 expression. Experimental hematology. 2019;69:27–36.

23. Boutzen H, Chan-Seng-Yue M, Murison A, Mbong N, Wagenblast E, Arlidge C, et al. A primary patient-derived model for investigating functional heterogeneity within the human Leukemic Stem Cell Compartment. bioRxiv. 2022:2022.03.01.482535.

24. Boutzen H, Murison A, Oriecuia A, Bansal S, Arlidge C, Wang JCY, et al. Identification of leukemia stem cell subsets with distinct transcriptional, epigenetic and functional properties. Leukemia. 2024;38(10):2090–101.

25. Boitano AE, Wang J, Romeo R, Bouchez LC, Parker AE, Sutton SE, et al. Aryl hydrocarbon receptor antagonists promote the expansion of human hematopoietic stem cells. Science (New York, NY). 2010;329(5997):1345–8.

26. Pabst C, Krosl J, Fares I, Boucher G, Ruel R, Marinier A, et al. Identification of small molecules that support human leukemia stem cell activity ex vivo. Nature methods. 2014;11(4):436–42.

27. Mistry H, Hsieh G, Buhrlage SJ, Huang M, Park E, Cuny GD, et al. Small-molecule inhibitors of USP1 target ID1 degradation in leukemic cells. Molecular cancer therapeutics. 2013;12(12):2651–62.

28. Arkinson C, Chaugule VK, Toth R, Walden H. Specificity for deubiquitination of monoubiquitinated FANCD2 is driven by the N-terminus of USP1. Life Science Alliance. 2018;1(5):e201800162.

29. Burkhart DL, Sage J. Cellular mechanisms of tumour suppression by the retinoblastoma gene. Nature Reviews Cancer. 2008;8(9):671–82.

30. Safa-Tahar-Henni S, Páez Martinez K, Gress V, Esparza N, Roques É, Bonnet-Magnaval F, et al. Comparative small molecule screening of primary human acute leukemias, engineered human leukemia and leukemia cell lines. Leukemia. 2025;39(1):29–41.

31. Guzman ML, Rossi RM, Neelakantan S, Li X, Corbett CA, Hassane DC, et al. An orally bioavailable parthenolide analog selectively eradicates acute myelogenous leukemia stem and progenitor cells. Blood. 2007;110(13):4427–35.

32. Flores-Lopez G, Moreno-Lorenzana D, Ayala-Sanchez M, Aviles-Vazquez S, Torres-Martinez H, Crooks PA, et al. Parthenolide and DMAPT induce cell death in primitive CML cells through reactive oxygen species. J Cell Mol Med. 2018;22(10):4899–912.

33. Zhou X, Moreira S, Restelli C, Wang H, Jahangiri S, Aryal S, et al. Activation of a nongenetic AHR-ELMSAN1 axis optimizes BET-targeting therapy and suppresses leukemia stem cells in preclinical models. Science Translational Medicine. 2025;17(810):eadn5400.

34. Williams SA, Maecker HL, French DM, Liu J, Gregg A, Silverstein LB, et al. USP1 deubiquitinates ID proteins to preserve a mesenchymal stem cell program in osteosarcoma. Cell. 2011;146(6):918–30.

35. García-Santisteban I, Peters GJ, Giovannetti E, Rodríguez JA. USP1 deubiquitinase: cellular functions, regulatory mechanisms and emerging potential as target in cancer therapy. Mol Cancer. 2013;12(1):91.

36. O’Brien CA, Kreso A, Ryan P, Hermans KG, Gibson L, Wang Y, et al. ID1 and ID3 regulate the self-renewal capacity of human colon cancer-initiating cells through p21. Cancer cell. 2012;21(6):777–92.

37. Tang R, Hirsch P, Fava F, Lapusan S, Marzac C, Teyssandier I, et al. High Id1 expression is associated with poor prognosis in 237 patients with acute myeloid leukemia. Blood. 2009;114(14):2993–3000.

38. Suh HC, Leeanansaksiri W, Ji M, Klarmann KD, Renn K, Gooya J, et al. Id1 immortalizes hematopoietic progenitors in vitro and promotes a myeloproliferative disease in vivo. Oncogene. 2008;27(42):5612–23.

39. Sonego M, Pellarin I, Costa A, Vinciguerra GLR, Coan M, Kraut A, et al. USP1 links platinum resistance to cancer cell dissemination by regulating Snail stability. Science Advances. 2019;5(5):eaav3235.

40. Liu D, Li Q, Zang Y, Li X, Li Z, Zhang P, et al. USP1 modulates hepatocellular carcinoma progression via the Hippo/TAZ axis. Cell Death & Disease. 2023;14(4):264.

41. Rennie ML, Gundogdu M, Arkinson C, Liness S, Frame S, Walden H. Structural and Biochemical Insights into the Mechanism of Action of the Clinical USP1 Inhibitor, KSQ-4279. Journal of medicinal chemistry. 2024;67(17):15557–68.

