## Supplemental Table Titles and Legends for "Discovery of compounds targeting human acute myeloid leukemia stem cells via a novel high-throughput screen"

### Supplemental Table Legends

**Table S1: Overview of the cell lines, patient samples and models screened in the high-throughput assay.** Description of the cell lines, patient samples and models by ID, Type, Source, Fusion/Oncogene, Subtype and Karyotype.

**Table S2: Overview of the compound library screened in the high-throughput assay.** Description of the library containing non-FDA/FDA-approved compounds, their companies and structurally diverse molecules synthesized by the IRIC medicinal chemistry platform. These are broken down into three categories to illustrate the total HTS drugs, 93 hits identified from primary screen and 50 confirmed compounds from dose response.

**Table S3: Description of non-FDA/FDA-approved inhibitors from APExBIO library.** Details on 1 960 compounds tested, including compound names, known targets and supplier IDs.

**Table S4: Non-FDA-approved inhibitors from selected companies.** Details on 4 410 non-FDA-approved inhibitors from commercial vendors.

**Table S5: List of identified inhibitors and activators from primary screen.** Contains list of 61 inhibitors and 32 activators identified with data on their inhibition or expansion on $CD34^+$ LSC-enriched OCI-AML-8227 cells, toxicity profiles on $CD34^+$ cord blood cells, drug name, supplier and library IDs.

**Table S6: Overlap of hits between $CD34^+$ LSC-enriched OCI-AML-8227 and AML cell lines.** Compounds active in ≥ 50% of seven AML cell lines (> 70% inhibition) with minimal toxicity (< 30% inhibition in both cord blood controls) were considered "cell line hits," yielding 66 compounds. Seventeen overlapped with the 61 anti-LSC candidates identified in OCI-AML-8227 $CD34^+$ cells, while 44 were unique to the LSC-enriched screen.

**Table S7: Dose response validation of 93 hits from primary screen.** Details of the 93 hits that were identified from primary screen and were validated in a two-fold 20-point dose response on the $CD34^+$ LSC-enriched fraction of OCI-AML-8227. Includes compound names, ID, Company, $IC_{50}$, graphical results, $R^2$ fit, max percent inhibition, minimum percent inhibition, separated based on inhibitors, activators and refined hits from secondary validation.

**Table S8: Dose response and $LC_{50}$ values of tertiary validated candidates.** Contains detailed results of the dose-response analysis of the 33 candidate compounds tested on OCI-AML-8227 using flow cytometry. Includes $LC_{50}$ values for total live cells, $CD34^-$ (total blasts), $CD34^+$ (total LSC-enriched fraction), $CD34^+CD38^+$ (LSC progenitor-enriched fraction), and $CD34^+CD38^-$ (LSC-enriched fraction). Classification of inhibitors is also listed.

**Table S9: Comparison of 20 candidates in LSC enriched models.** Comparison of $LC_{50}$ between both LSC models, OCI-AML-8227 and OCI-AML-20. Details on toxicity to OP9 stroma, supplier and compound ID are listed. Top five candidates that worked well in both models are highlighted.

**Table S10: BCLQ patient data and response classification to candidate compounds.** Summary of patient data for nine primary AML patient samples with OCI-AML-8227; primary AML ID, classification, tissue type, sampling status, age at diagnosis, gender, FAB classification, blasts (%), white blood cell count, karyotype, cytogenetic risk group, AML type, mutational status and responder status to corresponding compounds.

**Table S11: Differentially expressed genes following SJB2-043 24 hr treatment in OCI-AML-8227 cells from scRNA-seq.** Gene list (normalized to DMSO untreated) showing $log_2FC$, p-values, FDR, pct.1, pct.2, comparison, ranked list (-log10(pvalue)*log2FC) and cell subset.

**Table S12: Differentially expressed genes following BIO-acetoxime 24 hr treatment in OCI-AML-8227 cells from scRNA-seq.** Gene list (normalized to DMSO untreated) showing $\log_2$FC, p-values, FDR, pct.1, pct.2, comparison, ranked list (-log10(pvalue)*log2FC) and cell subset.

**Table S13: Differentially expressed genes following Uxxxxx03 24 hr treatment in OCI-AML-8227 cells from scRNA-seq.** Gene list (normalized to DMSO untreated) showing $\log_2$FC, p-values, FDR, pct.1, pct.2, comparison, ranked list (-$\log_{10}$(pvalue)*$\log_2$FC) and cell subset.

**Table S14: GSEA results for SJB2-043 24 hr treatment compared to DMSO control for HALLMARK and KEGG pathways.** GSEA name of the signature, size, enrichment score (ES), normalized enrichment score (NES), normalized p-value (NOM p-val), FDR, FEWER p-value, rank at max and leading-edge percentages are shown for positively and negatively enriched signatures. Significant signatures are boxed (black) and were based on NOM p-value < 0.05 and FDR < 0.25.

**Table S15: GSEA results for BIO-acetoxime 24 hr treatment compared to DMSO control for HALLMARK and KEGG pathways.** GSEA name of the signature, size, enrichment score (ES), normalized enrichment score (NES), normalized p-value (NOM p-val), FDR, FEWER p-value, rank at max and leading-edge percentages are shown for positively and negatively enriched signatures. Significant signatures are boxed (black) and were based on NOM p-value < 0.05 and FDR < 0.25.

**Table S16: GSEA results for UMxxxxx03 treatment compared to DMSO control for HALLMARK and KEGG pathways.** GSEA name of the signature, size, enrichment score (ES), normalized enrichment score (NES), normalized p-value (NOM p-val), FDR, FEWER p-value, rank at max and leading-edge percentages are shown for positively and negatively enriched signatures. Significant signatures are boxed (black) and were based on NOM p-value $< 0.05$ and FDR $< 0.25$.
