## Supplemental Figures for "Discovery of compounds targeting human acute myeloid leukemia stem cells via a novel high-throughput screen"

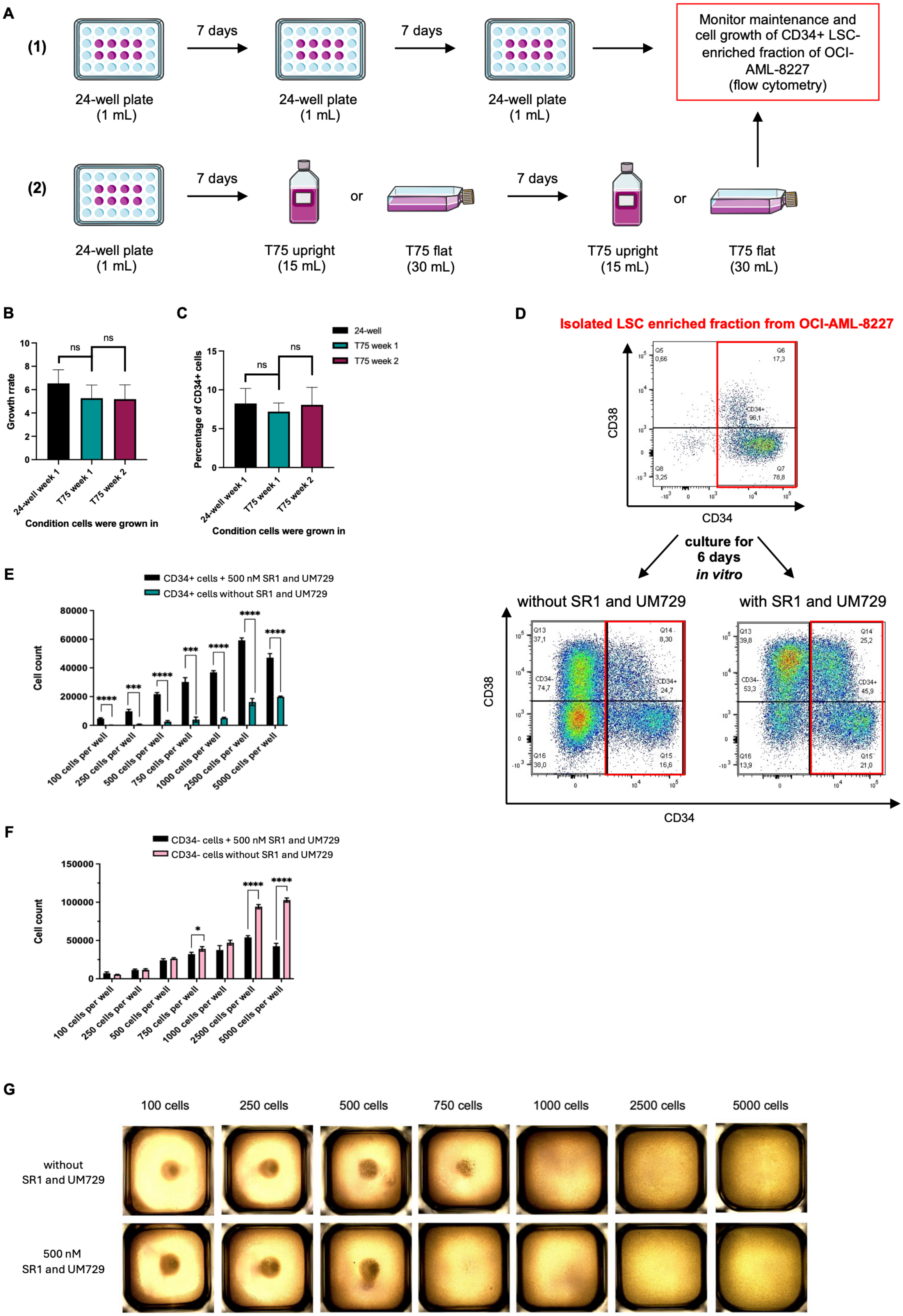
A
(1)
7 days
7 days
24-well plate
(1 mL)
24-well plate
(1 mL)
24-well plate
(1 mL)
Monitor maintenance and cell growth of CD34+ LSC-enriched fraction of OCI-AML-8227
(flow cytometry)
(2)
7 days
or
7 days
or
24-well plate
(1 mL)
T75 upright
(15 mL)
T75 flat
(30 mL)
T75 upright
(15 mL)
T75 flat
(30 mL)
B
Growth rrate
ns
ns
24-well week 1
T75 week 1
T75 week 2
Condition cells were grown in
C
24-well
T75 week 1
T75 week 2
Percentage of CD34+ cells
ns
ns
24-well week 1
T75 week 1
T75 week 2
Condition cells were grown in
D
Isolated LSC enriched fraction from OCI-AML-8227
CD38
CD34
culture for
6 days
in vitro
without SR1 and UM729
with SR1 and UM729
CD38
CD34
E
CD34+ cells + 500 nM SR1 and UM729
CD34+ cells without SR1 and UM729
Cell count
100 cells per well
250 cells per well
500 cells per well
750 cells per well
1000 cells per well
2500 cells per well
5000 cells per well
F
CD34- cells + 500 nM SR1 and UM729
CD34- cells without SR1 and UM729
Cell count
100 cells per well
250 cells per well
500 cells per well
750 cells per well
1000 cells per well
2500 cells per well
5000 cells per well
G
100 cells
250 cells
500 cells
750 cells
1000 cells
2500 cells
5000 cells
without
SR1 and UM729
500 nM
SR1 and UM729

**H**

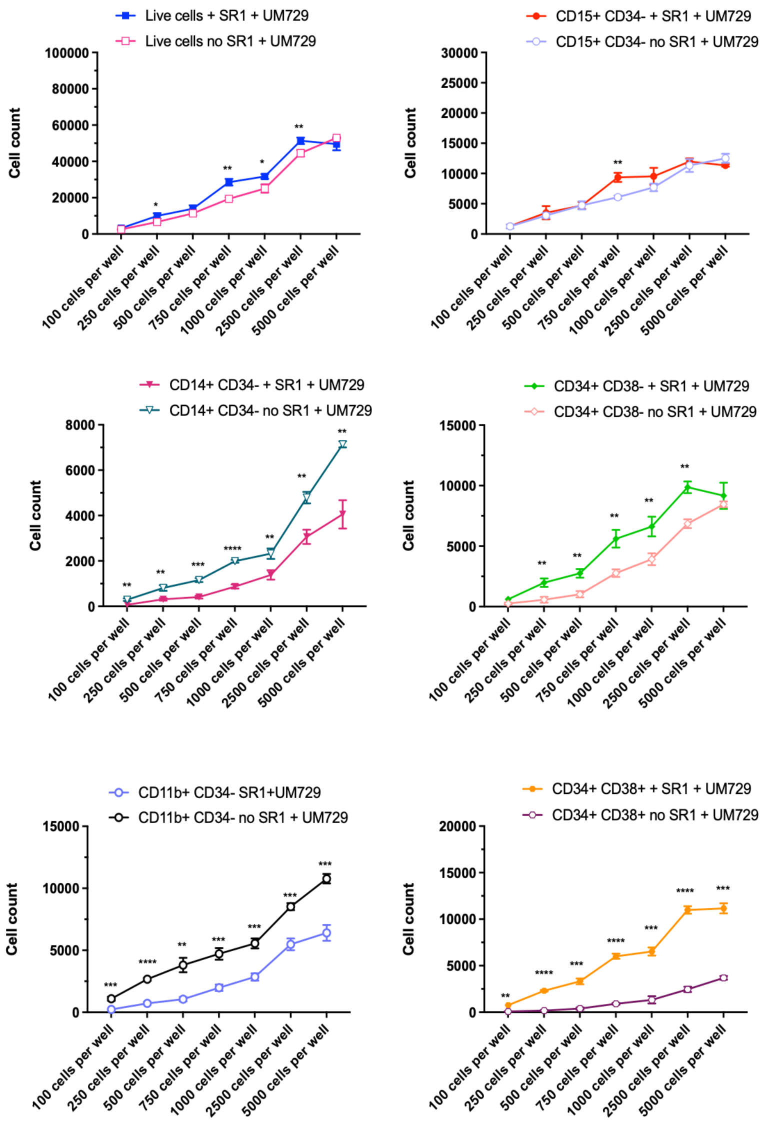


**Figure S1: Optimization of culture conditions and use of differentiation inhibitors for high-throughput screening in the LSC-enriched population of OCI-AML-8227. (A)** Schematic of culture format comparison: OCI-AML-8227 cells grown in 24-well plates (1 mL) or scaled to T75 flasks (upright 15 mL, flat 30 mL) for two weeks. Orientation and volumes were chosen based on the oxygen to $CO_2$ consumption ratio. Cells were monitored by flow cytometry. **(B-C)** Growth rate **(B)** and percentage of viable $CD34^+$ cells (normalized to DMSO) **(C)** between T75 flasks and 24-

well plates over two weeks of culture, demonstrating no significant difference between the two culturing methods. Mean ± s.d., unpaired t-test. Representative one of $n = 1$ biological replicate. **(D)** Flow cytometry profile (CD38 vs CD34) of the isolated $CD34^+$ LSC-enriched fraction of OCI-AML-8227 over six days with or without 500 nM SR1 and UM729 (5 000 cells initially plated/well). **(E-F)** Cell count of $CD34^+$ cells **(E)** and $CD34^-$ blasts **(F)** at different seeding densities. Mean ± s.d. Representation one of $n = 2$ biological replicates. **(G)** Representative images of OCI-AML-8227 seeded at 100–5 000 $CD34^+$ cells/well in 384-well plates with or without 500 nM SR1 and UM729. Photos were taken with Invitrogen™ EVOS™ XL Core Imaging System. **(H)** Expanded analysis of additional phenotypic subsets ($CD15^+CD34^-$, $CD11b^+CD34^-$, $CD14^+CD34^-$ blasts), demonstrating reduced differentiation and increased $CD34^+$ cell numbers ($CD34^+CD38^+$ leukemic progenitors and $CD34^+CD38^-$ LSC-enriched) across all densities with SR1 and UM729. Data are mean ± s.d.; $^*p < 0.05$, $^{**}p < 0.01$, $^{***}p < 0.001$, $^{****}p < 0.0001$ (unpaired t-test unless otherwise stated). Representation one of $n = 2$ biological replicates.

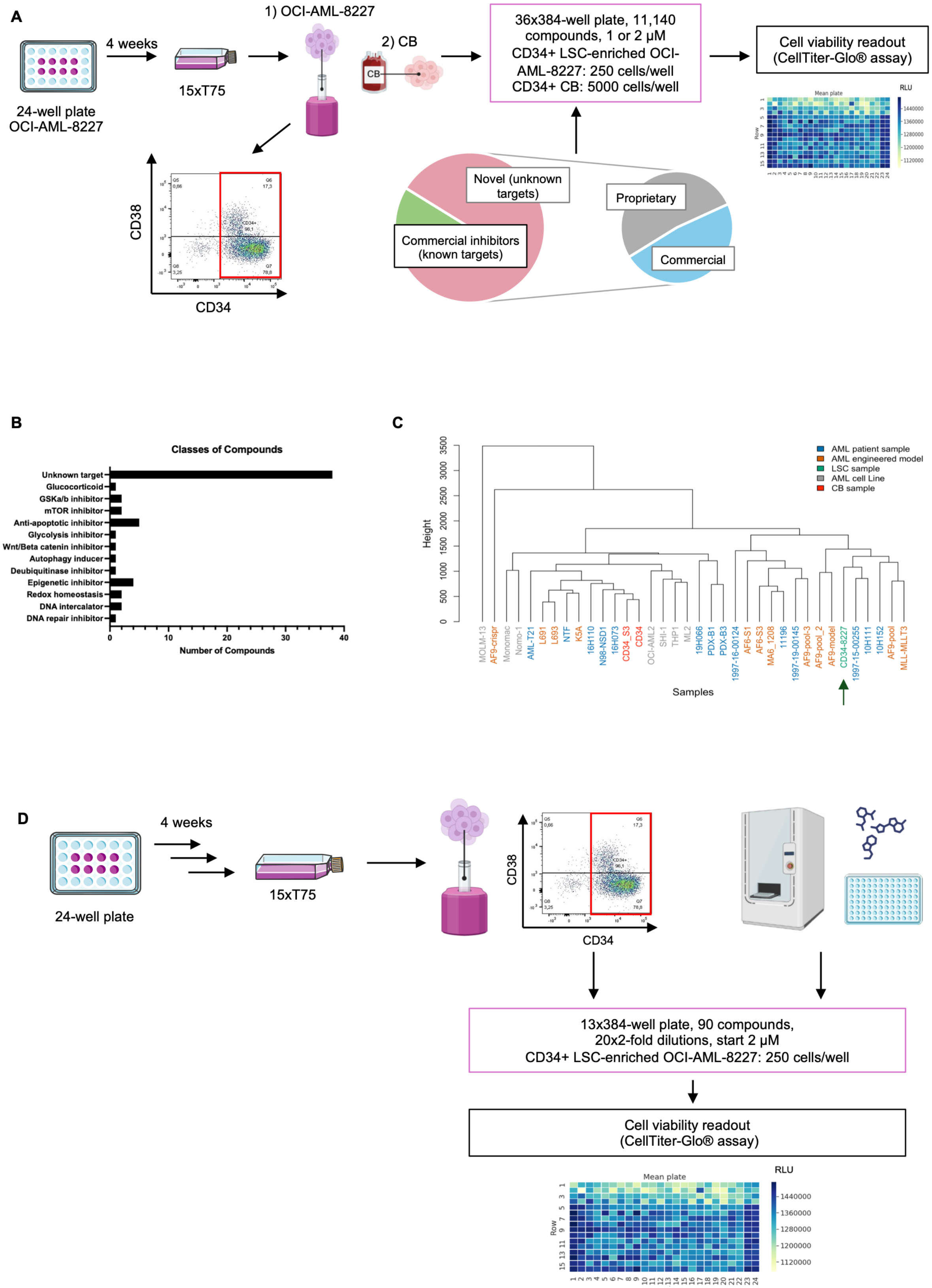
A
4 weeks
1) OCI-AML-8227
2) CB
CB
24-well plate
OCI-AML-8227
15xT75
36x384-well plate, 11,140 compounds, 1 or 2 µM
CD34+ LSC-enriched OCI-AML-8227: 250 cells/well
CD34+ CB: 5000 cells/well
Cell viability readout (CellTiter-Glo® assay)
Mean plate
RLU
Row
CD38
CD34
Novel (unknown targets)
Commercial inhibitors (known targets)
Proprietary
Commercial
B
Classes of Compounds
Unknown target
Glucocorticoid
GSKa/b inhibitor
mTOR inhibitor
Anti-apoptotic inhibitor
Glycolysis inhibitor
Wnt/Beta catenin inhibitor
Autophagy inducer
Deubiquitinase inhibitor
Epigenetic inhibitor
Redox homeostasis
DNA intercalator
DNA repair inhibitor
Number of Compounds
C
Height
AML patient sample
AML engineered model
LSC sample
AML cell Line
CB sample
MOLM-13
AF9-crispr
Monomac
Nomo-1
AML-T21
L691
L693
NTF
K5A
16H110
N98-NSD1
16H073
CD34_S3
CD34
OCI-AML2
SHI-1
THP1
ML2
19H066
PDX-B1
PDX-B3
1997-16-00124
AF6-S1
AF6-S3
MA6_1208
11196
1997-19-00145
AF9-pool-3
AF9-pool_2
AF9-model
CD34-8227
1997-15-00255
10H111
10H152
AF9-pool
MLL-MLLT3
Samples
D
4 weeks
24-well plate
15xT75
CD38
CD34
13x384-well plate, 90 compounds,
20x2-fold dilutions, start 2 µM
CD34+ LSC-enriched OCI-AML-8227: 250 cells/well
Cell viability readout
(CellTiter-Glo® assay)
Mean plate
RLU
Row

**Figure S2: Overview of high-throughput screening workflow, hit classification and validation strategy in CD34$^+$ LSC-enriched fraction of OCI-AML-8227.**

**(A)** Schematic of HTS: 11 142 compounds were tested, comprising 1 960 FDA-approved or non-approved molecules with known targets and 9 182 non-approved compounds of unknown target. CD34$^+$ LSC-enriched OCI-AML-8227 cells and CD34$^+$ CB controls were treated with 1 or 2 μM compounds. *N* = 1 biological replicate. **(B)** Target classification of 61 inhibitors (23 known targets, 38 unknown) and 32 activators. **(C)** Unsupervised hierarchical clustering of drug sensitivity across 34 AML patient samples, AML models, and CB. Inclusion criteria: >70% inhibition in at least one AML sample and <30% inhibition in two CB samples. CD34$^+$ OCI-AML-8227 cells is marked with green arrow. **(D)** Schematic of secondary validation: 101 compounds (93 primary screen hits and 8 controls) were tested in a dose response assay on CD34$^+$ LSC-enriched OCI-AML-8227 cells (250 cells/well, duplicates) supplemented with 500 nM SR1 and UM729. *N* = 1 biological replicate.

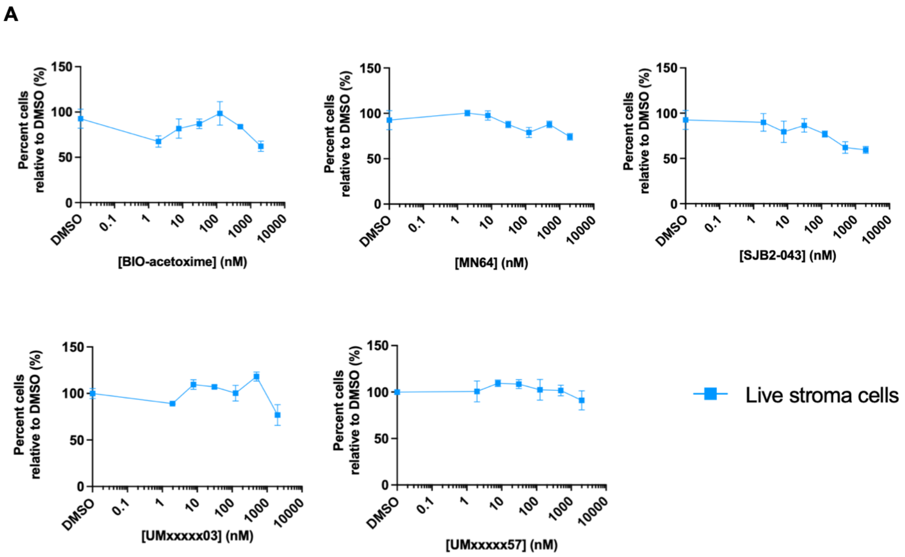


**Figure S3: Lead compounds active in OCI-AML-20 show minimal toxicity to stromal cells.** **(A)** OP9 stromal cells were treated for six days with the indicated compounds at the same doses used in OCI-AML-20 assays. Viability was measured by SYTOX exclusion. All five lead inhibitors that produced ≥50% reduction in $CD34^+$ LSC-enriched OCI-AML-20 fractions showed minimal stromal toxicity, confirming selectivity for leukemic over niche cells. Data are mean ± s.d. Representative one of $n$ = 3 biological replicates.

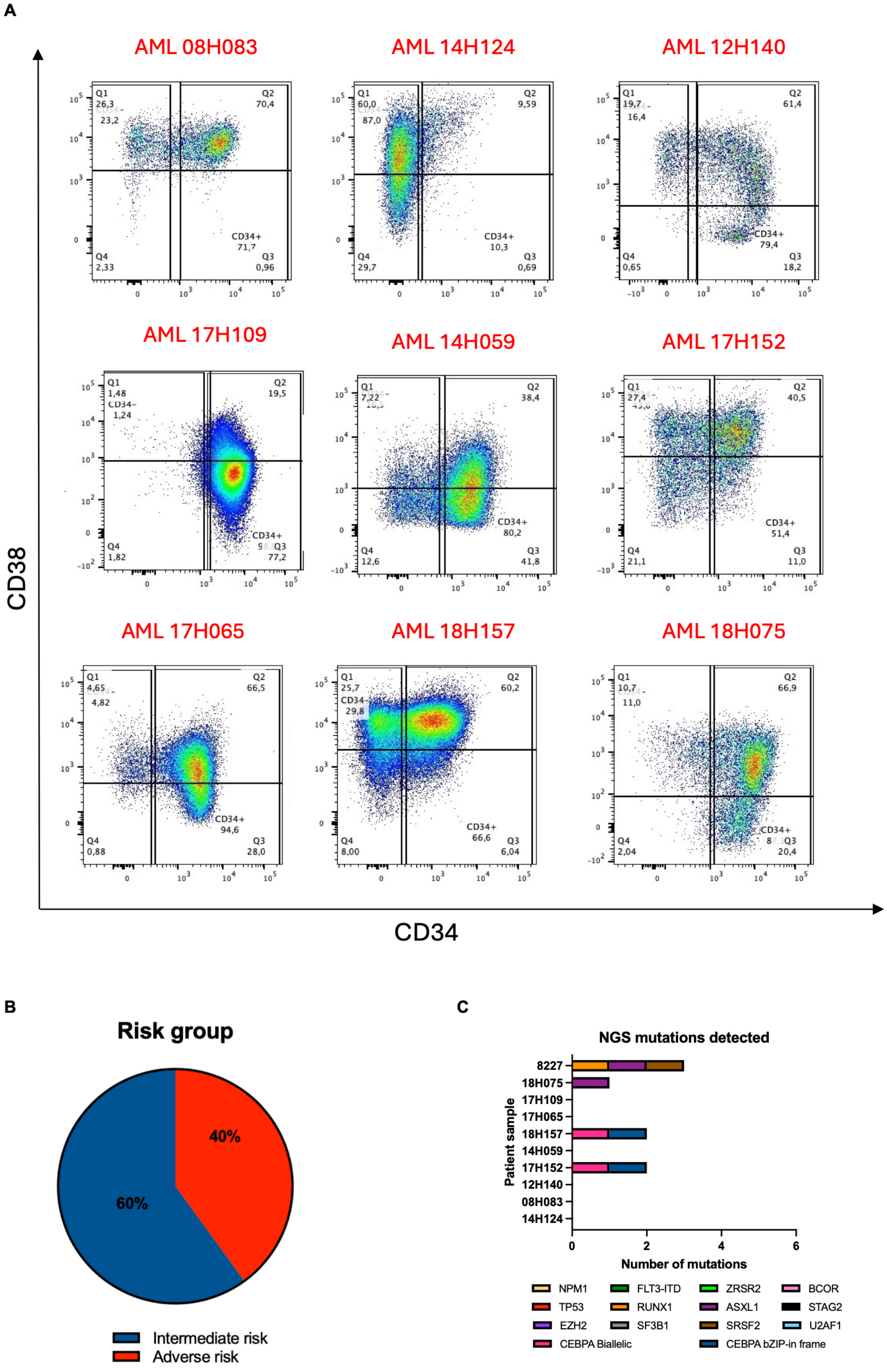
A
AML 08H083
AML 14H124
AML 12H140
AML 17H109
AML 14H059
AML 17H152
AML 17H065
AML 18H157
AML 18H075
CD38
CD34
B
Risk group
40%
60%
Intermediate risk
Adverse risk
C
NGS mutations detected
Patient sample
8227
18H075
17H109
17H065
18H157
14H059
17H152
12H140
08H083
14H124
Number of mutations
NPM1
FLT3-ITD
ZRSR2
BCOR
TP53
RUNX1
ASXL1
STAG2
EZH2
SF3B1
SRSF2
U2AF1
CEBPA Biallelic
CEBPA bZIP-in frame

**Figure S4: Flow cytometry profile for nine primary AML patient samples from BCLQ. (A)** Flow cytometry profile of all nine primary AML patient samples for CD38 vs CD34, illustrating gating for: $CD34^{-}CD38^{-}$, $CD34^{-}CD38^{+}$, $CD34^{+}CD38^{+}$ and $CD34^{+}CD38^{-}$ populations. Percentages in each gate corresponding to live populations. Cells were cultured in the presence of 500 nM SR1 and UM729 to maintain $CD34^{+}$ LSC-enriched populations. **(B)** Risk group distribution of ten primary AML (including OCI-AML-8227) (blue = intermediate risk (60%), red = adverse risk (40%)). **(C)** Mutational landscape confirmed by next-generation sequencing. $N = 1$ biological replicate.

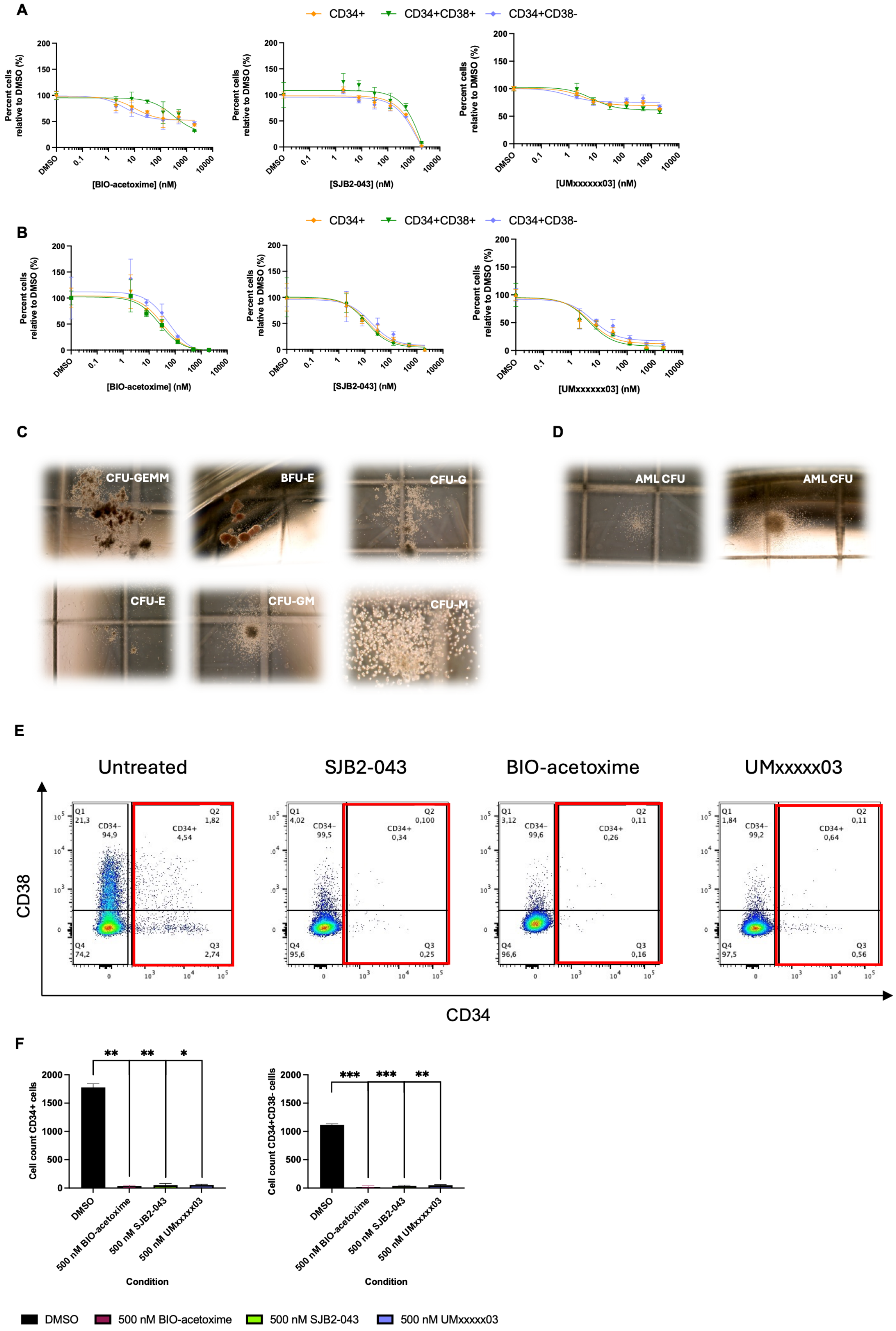
A
CD34+
CD34+CD38+
CD34+CD38-
Percent cells relative to DMSO (%)
[BIO-acetoxime] (nM)
[SJB2-043] (nM)
[UMxxxxxx03] (nM)
DMSO
0.1
1
10
100
1000
10000
B
CD34+
CD34+CD38+
CD34+CD38-
Percent cells relative to DMSO (%)
[BIO-acetoxime] (nM)
[SJB2-043] (nM)
[UMxxxxxx03] (nM)
C
CFU-GEMM
BFU-E
CFU-G
CFU-E
CFU-GM
CFU-M
D
AML CFU
AML CFU
E
Untreated
SJB2-043
BIO-acetoxime
UMxxxxx03
CD38
CD34
Q1 21,3
CD34- 94,9
Q2 1,82
CD34+ 4,54
Q4 74,2
Q3 2,74
Q1 4,02
CD34- 99,5
Q2 0,100
CD34+ 0,34
Q4 95,6
Q3 0,25
Q1 3,12
CD34- 99,6
Q2 0,11
CD34+ 0,26
Q4 96,6
Q3 0,16
Q1 1,84
CD34- 99,2
Q2 0,11
CD34+ 0,64
Q4 97,5
Q3 0,56
F
Cell count CD34+ cells
Cell count CD34+CD38- cellls
**
**
*
***
***
**
DMSO
500 nM BIO-acetoxime
500 nM SJB2-043
500 nM UMxxxxx03
Condition
DMSO
500 nM BIO-acetoxime
500 nM SJB2-043
500 nM UMxxxxx03

**Figure S5: Lead compounds show selective activity against LSC-enriched cells over healthy HSPCs and reduce leukemic progenitor and LSC function while sparing normal hematopoietic progenitors. (A–B)** $CD34^+$ cord blood (HSPCs) **(A)** and $CD34^+$ LSC-enriched OCI-AML-8227 cells **(B)** were treated for six days with increasing doses of BIO-acetoxime, SJB2-043, or UMxxxxx03. Subpopulations were analyzed by flow cytometry to determine $LC_{50}$ values. Mild toxicity to HSPCs was observed only at higher doses (>500 nM), with $LC_{50}$ values ~20–100× higher than for LSC-enriched cells, indicating preferential targeting of LSCs over healthy hematopoietic cells. Data are mean ± s.d. Representative one of $n = 3$ biological replicates. **(C-D)** Representative images (one of $n = 3$ biological replicates) of colony-forming unit (CFU) assays from CB $CD34^+$ cells **(C)** and OCI-AML-8227 **(D)** after six-day treatment with BIO-acetoxime, SJB2-043, or UMxxxxx03, followed by 12 days in methylcellulose. Colonies scored as CFU-E, BFU-E, CFU-GM, CFU-G, CFU-M, or CFU-GEMM for CB **(C)** and AML-CFU for OCI-AML-8227 **(D)**. Photos were taken with Invitrogen™ EVOS™ XL Core Imaging System. **(E)** Flow cytometry plots showing $CD34^+$ cell percentages in OCI-AML-8227 after seven-day treatment with BIO-acetoxime, SJB2-043, or UMxxxxx03, prior to intrafemoral injection into NSG-S mice. **(F)** Absolute counts of $CD34^+$ cells (left) and $CD34^+CD38^-$ cells (right) prior to injection. $N = 2$ biological replicates, $n = 10$ mice per group. Unpaired t-test $p < 0.05$, $^{**}p < 0.01$, $^{***}p < 0.001$, $^{****}p < 0.0001$.

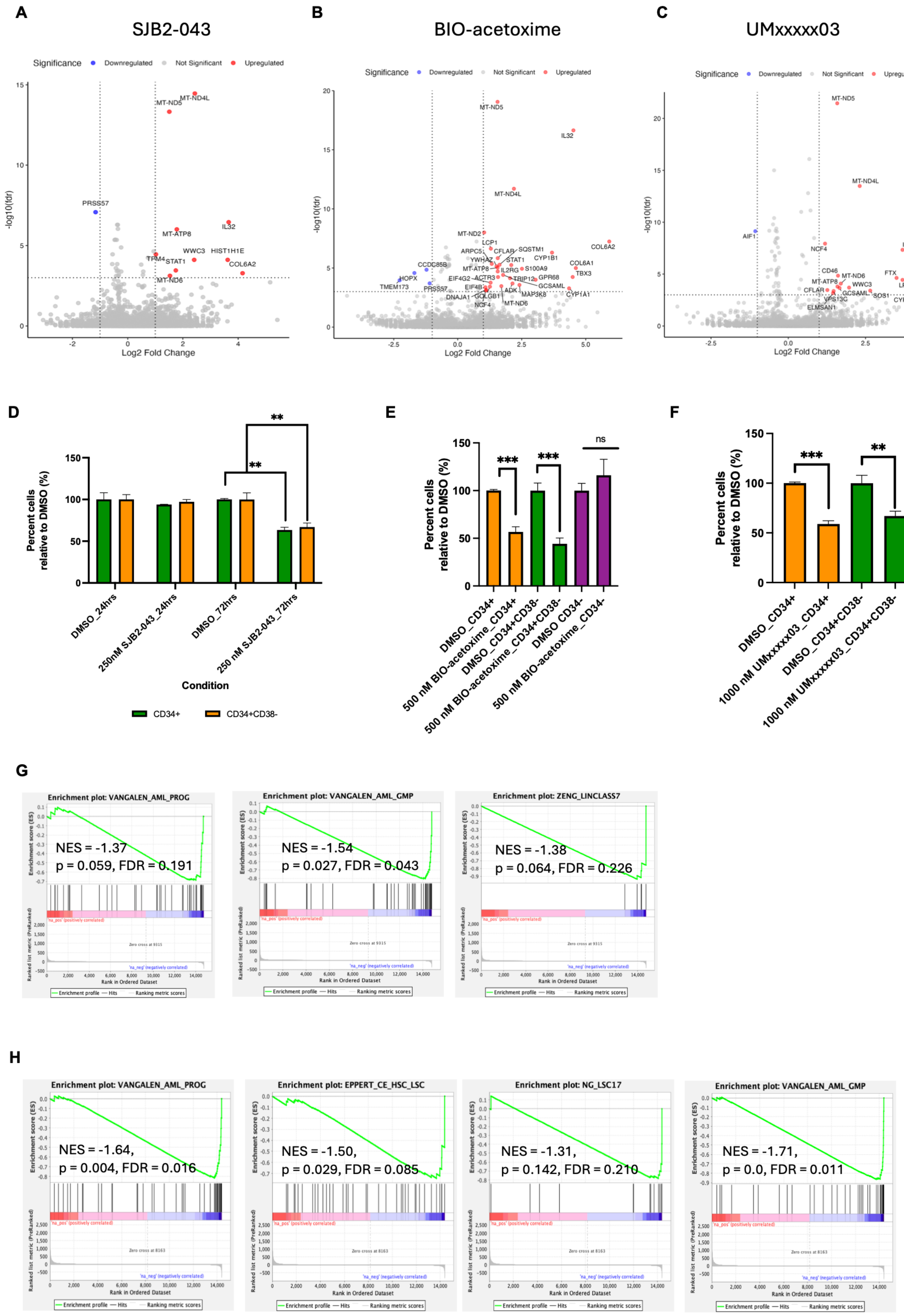
A
SJB2-043
Significance Downregulated Not Significant Upregulated
-log10(fdr)
Log2 Fold Change
MT-ND4L
MT-ND5
PRSS57
IL32
MT-ATP8
WWC3
HIST1H1E
TPM4
STAT1
COL6A2
MT-ND6
B
BIO-acetoxime
Significance Downregulated Not Significant Upregulated
-log10(fdr)
Log2 Fold Change
MT-ND5
IL32
MT-ND4L
MT-ND2
LCP1
COL6A2
ARPC5
CFLAR
SQSTM1
YWHAZ
STAT1
CYP1B1
CCDC85B
MT-ATP8
IL2RG
S100A9
COL6A1
TBX3
HOPX
EIF4G2
ACTR3
TRIP12
GPR68
TMEM173
PRSS57
EIF4B
ADK
GCSAML
DNAJA1
GOLGB1
MAP3K8
CYP1A1
NCF4
MT-ND6
C
UMxxxxx03
Significance Downregulated Not Significant Upregulated
-log10(fdr)
Log2 Fold Change
MT-ND5
MT-ND4L
AIF1
NCF4
IL32
TBX3
CD46
MT-ND6
FTX
MT-ATP8
WWC3
LPAR6
CFLAR
GCSAML
SOS1
CYP1A1
VPS13C
ELMSAN1
D
Percent cells relative to DMSO (%)
Condition
DMSO_24hrs
250nM SJB2-043_24hrs
DMSO_72hrs
250 nM SJB2-043_72hrs
CD34+
CD34+CD38-
**
**
E
Percent cells relative to DMSO (%)
DMSO_CD34+
500 nM BIO-acetoxime_CD34+
DMSO_CD34+CD38-
500 nM BIO-acetoxime_CD34+CD38-
DMSO CD34-
500 nM BIO-acetoxime_CD34-
***
***
ns
F
Percent cells relative to DMSO (%)
DMSO_CD34+
1000 nM UMxxxxx03_CD34+
DMSO_CD34+CD38-
1000 nM UMxxxxx03_CD34+CD38-
***
**
G
Enrichment plot: VANGALEN_AML_PROG
NES = -1.37
p = 0.059, FDR = 0.191
Enrichment plot: VANGALEN_AML_GMP
NES = -1.54
p = 0.027, FDR = 0.043
Enrichment plot: ZENG_LINCLASS7
NES = -1.38
p = 0.064, FDR = 0.226
Enrichment score (ES)
Ranked list metric (PreRanked)
Rank in Ordered Dataset
Enrichment profile — Hits Ranking metric scores
H
Enrichment plot: VANGALEN_AML_PROG
NES = -1.64,
p = 0.004, FDR = 0.016
Enrichment plot: EPPERT_CE_HSC_LSC
NES = -1.50,
p = 0.029, FDR = 0.085
Enrichment plot: NG_LSC17
NES = -1.31,
p = 0.142, FDR = 0.210
Enrichment plot: VANGALEN_AML_GMP
NES = -1.71,
p = 0.0, FDR = 0.011
Enrichment score (ES)
Ranked list metric (PreRanked)
Rank in Ordered Dataset
Enrichment profile — Hits Ranking metric scores

113

I

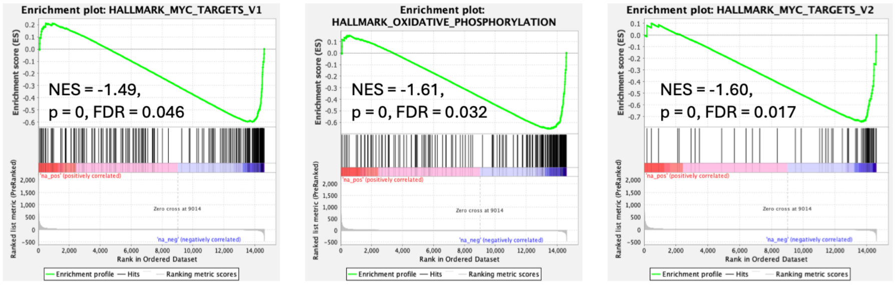


**Figure S6: Shared stress-response pathways but distinct functional outcomes following treatment with lead compounds.** **(A–C)** scRNA-seq volcano plots of differentially expressed genes (DEGs) showing significantly upregulated (red) and downregulated (blue) genes in LSC cluster after 24 h treatment with 250 nM SJB2-043 **(A)**, 500 nM BIO-acetoxime **(B)**, or 1 000 nM UMxxxxx03 **(C)**. Cut-off of $|\log_2 FC>1|$ and FDR < 0.001 for significant genes. Genes in grey are nonsignificant. $N$ = 1 biological replicate. **(D–F)** Flow cytometry quantification of viable $CD34^+$ and $CD34^+CD38^-$ populations after 24 or 72 h treatment, relative to DMSO control, showing LSC depletion with SJB2-043 (24 and 72 hr) **(D),** BIO-acetoxime (72 hr) **(E)**, and UMxxxxx03 (72 hr) **(F)**. Data are mean ± s.d.; $*p < 0.05$, $**p < 0.01$, $***p < 0.001$, $****p < 0.0001$ (unpaired t-test unless otherwise stated). Non-significant: ns. $N$ = 1 biological replicate. **(G-I)** GSEA plots of selected stemness- and progenitor-associated signatures in the multi-lineage GMP cluster after treatment with SJB2-043 **(G)**, BIO-acetoxime **(H)**, or UMxxxxx03 **(I)**, showing consistent negative enrichment of LSC-associated programs. $N$ = 1 biological replicate.
