## Supplemental Methods for "Discovery of compounds targeting human acute myeloid leukemia stem cells via a novel high-throughput screen"

### Supplemental Materials, Methods, Figures and Tables

#### Optimizing cell culturing conditions for OCI-AML-8227 expansion

OCI-AML-8227 cells were grown in different vessels for two weeks: 24-well plates (Falcon®) and T75 cell culture flasks (different orientations; 30 mL flat and 15 mL upright) (Life Technologies) in StemSpan™ serum-free media II (SFEM II) (STEMCELL Technologies) containing penicillin-streptomycin (Life Technologies, 1:1000, pen/strep) and growth factors: 10 ng/mL interleukin 3 (IL-3), IL-6 and granulocyte colony-stimulating factor (G-CSF), 25 ng/mL thrombopoietin (TPO), 50 ng/mL stem cell factor (SCF) and FLT3 ligand (FLT3L) (Life Technologies). Cell viability and maintenance of LSC phenotype ($CD34^+$) was assessed using antibodies for cell surface markers CD34 (APC, RRID:AB_ AB_1877153), CD38 (PE, RRID:AB_2561900) and CD15 (FITC, RRID:AB_314196) (Biolegend) and counting by hemacytometer with trypan blue staining. SYTOX™ Blue Dead Cell Stain was used for cell viability for flow cytometry analysis (Life Technologies). Flow cytometry was performed using a LSR Fortessa (BD Bioscience).

#### Optimizing isolation/enrichment and maintenance of OCI-AML-8227 $CD34^+$ cells

Cells were enriched using SFEM II and the EasySep™ Magnet and EasySep™ CD34 Positive Selection Kit II protocol (Catalog #17856, STEMCELL Technologies). Cells were then washed in SFEM II and counted using a hemocytometer to determine final yield. Cells were stained with previously mentioned antibodies and flow cytometry was performed using a LSRFortessa (BD Biosciences). Isolated $CD34^+$ cells were plated at various cell concentrations: 100, 250, 500, 750, 1000, 2500 and 5,000 cells/well in SFEM II media containing pen/strep and all six growth factors previously mentioned in a 384-well plate (Corning® catalog #3570). Two conditions were tested:

1) StemRegenin1 (SR1, Cat. No.: #72342) and UM729 (Cat. No.: #72332 (STEMCELL Technologies) at 500 nM and 2) without these compounds. After six days, photos of each cell concentration were taken to assess cell density and confluency using an Invitrogen™ EVOS™ XL Core Imaging System. Cells were collected, washed with 2% Cosmic Calf Serum (CCS) (Fisher Scientific) in 1X pH 7.4 phosphate buffer (PBS) (0) calcium chloride and magnesium chloride (Life Technologies) (2% CCS in PBS) and stained with cell surface markers previously mentioned. Cell count was assessed using flow cytometry (LSR Fortessa) fitted with a high-throughput samples (HTS) (BD Bioscience). The ratio between $CD34^{+}$ and $CD34^{-}$ cells with and without SR1 and UM729 were compared. Student unpaired t-test was performed to compare both conditions using GraphPad Prism v10 for [Windows/Mac].

**High-throughput screening of OCI-AML-8227, 34 primary patient samples, models and cell lines and cord blood.**

OCI-AML-8227 cells were expanded and enriched for $CD34^{+}$ cells using the EasySep™ CD34 Positive Selection Kit II (STEMCELL Technologies, #17856) and plated at 250 cells/well in StemSpan™ serum-free media II (SFEM II, Cat. No. #09655) (STEMCELL Technologies) supplemented with (Life Technologies, 1:1000, pen/strep) and growth factors: 10 ng/mL interleukin (IL3)-3, IL-6 and granulocyte colony-stimulating factor (G-CSF), 25 ng/mL thrombopoietin (TPO), 50 ng/mL stem cell factor (SCF) and FLT3 ligand (FLT3L) (Life Technologies). SR1 and UM729 were added at 500 nM throughout primary and secondary screening. Cells were dispensed into 36 × 384-well plates containing compounds using an Echo 555 acoustic dispenser and Multidrop Combi nL reagent dispenser. Compounds were tested at 1 µM (APExBIO, FDA-approved and commercial libraries) or 2 µM (University of Montreal and IRIC libraries), with a final DMSO concentration of 0.1%. The 11 142 compound collection comprised molecules from APExBIO (1 960 compounds), ChemBridge (2 572), Maybridge (1 463), SPECS (363), Cayman Chemicals (4), and proprietary libraries (4 772 compounds), including 3 471 synthesized by the IRIC Medicinal Chemistry Facility. After six days of incubation, viability was assessed using the CellTiter-Glo® Luminescent Assay (Promega). Primary screen normalization, quality control metrics, and hit selection criteria are described below.

To assess toxicity toward normal hematopoietic cells, $CD34^+$ cells were isolated from fresh human cord blood (<24 h post-collection; CHU Sainte-Justine) by magnetic selection and screened under identical conditions at 5,000 cells/well. All samples were obtained with informed consent under approved institutional protocols. For culturing and testing of primary patient samples, models and cell lines, see reference (1).

**Data analysis of high-throughput screen**

Percent inhibition scores were corrected for drift in DMSO controls and normalized as stated elsewhere (1):

$$Cscore = 100 - \left(\frac{X_i}{\overline{C_j}} \times 100\right)$$

Where $X_i$ is the luminescence value (in RLU) of well $i$ in row $j$ and $\overline{C_j}$ is the mean luminescence value of the control wells in row $j$. Quality control was performed as mentioned previously (1). Criteria for percent inhibition on $CD34^+$ 8227 cells and $CD34^+$ cord blood cells were compared to generate hits, as well as with the other AML models screened (*n*=33 samples). A total of 667 hits were generated where 574 were removed. Criteria for selecting hits were >70% inhibition on

$CD34^+$ OCI-AML-8227 cells and <30% on $CD34^+$ cord blood cells. This led to identifying 61 inhibitors and 32 activators (provided expansion of cells through differentiation).

To compare the response similarity of $CD34^+$ OCI-AML-8227 with other AML patient samples, models and cells, a heat map with unsupervised hierarchal clustering using R packages pheatmap (RRID:SCR_016418) and hclust (RRID:SCR_014673), a criterion of >70% on at least 1 AML sample and <30% on 2 $CD34^+$ cord bloods was used. To assess common hits with $CD34^+$ OCI-AML-8227 with 7 cell lines, a criterion of <30% on both cord bloods used and >70% inhibition on 4/7 cell lines. A Venn diagram created using R package VennDiagram (RRID:SCR_002414) was used to compare the common hits among the cell lines and $CD34^+$ OCI-AML-8227.

For secondary dose response screen, percent inhibition scores were normalized to DMSO controls, %Inh= 100 – ((RLU of sample) / (Average of RLU DMSO))*100. RLU is the luminescent in relative light units. Secondary hits were chosen based on best fit and $IC_{50}$ ($R^2$>0.70 and $IC_{50}$ < 1000 nM).

**Dose response screen on a second-LSC enriched model, OCI-AML-20**

OP9 stroma cells (5 000 per well) were plated in 96-well TC-treated flat bottom plates in Minimal Essential Media (MEM) Alpha (1X) + GlutaMAX™ (-nucleosides) (Fisher Scientific) with 20% heat inactivated fetal bovine serum (FBS) (Wisent), 55 µM b-mercaptoethanol (Fisher Scientific) and 0.01% penicillin-streptomycin (Life Technologies), 48 hours prior to seeding OCI-AML-20. OP9 media was then removed, wells were washed with Gibco™ PBS pH 7.4 (1X) (Fisher Scientific) and OCI-AML-20 (50 000 per well) were seeded on OP9 cells in IMDM + 2 mM L-glutamine + 25 mM HEPES (Fisher Scientific), 10% heat inactivated FBS, 55 µM bmercaptoethanol (Fisher Scientific), 0.01% penicillin-streptomycin and 20 ng/mL of granulocyte macrophage colony-stimulating factor (GM-CSF) (Life Technologies). For OP9 only control condition, the same procedure applies as above, but the OCI-AML-20 were not added to each well, only the media. OCI-AML-8227 cells were plated separately (50 000 per well) in corresponding media. The cells were treated with the 20 candidates at 6 different concentrations (DMSO 0.02%, 1.95 nM, 7.8 nM, 31 nM, 125 nM, 500 nM, 2 000 nM) in triplicates. Cells were incubated for six days. OCI-AML-20 and OP9 cells were then trypsinized, collected and washed in Gibco™ PBS pH 7.4 (1X). For OCI-AML-8227, cells were collected and washed with 2% CCS in PBS. Cells were then stained with antibodies to CD34 (APC, RRID:AB_1877153), CD38 (PE, RRID:AB_2561900), CD15 (FITC, RRID:AB_AB_314196), CD11b (BV650, RRID:AB_2563793) and CD45 (Alex Fluor®700, RRID:AB_2566372) (Biolegend) and viability dye SYTOX™ Blue Dead Cell Stain (Life Technologies). Flow cytometry was performed using an LSR Fortessa fitted with a high-throughput sampler (HTS). The $LC_{50}$ was determined for all OCI-AML-20 and -8227 populations using GraphPad Prism software nonlinear regression (curve fit) – dose-response – inhibition and the toxicity on OP9 stroma.

**Apoptosis assay**

OCI-AML-8227, OCI-AML-20, and OP9 (RRID: CVCL_4398) cells were treated for three days with candidate compounds at 0.05% DMSO, 2 000 nM, or 5 000 nM, with cytarabine included as a positive control for apoptosis. Cells were stained with antibodies to CD34 (APC, RRID:AB_ AB_1877153), CD38 (PE, RRID:AB_2561900), CD15 (FITC, RRID:AB_314196), CD11b (BV650, RRID: AB_2563793), and CD45 (AlexaFluor®700, RRID:AB_2566374), followed by Annexin V Binding Buffer (BioLegend, Cat. No.: #422201), Annexin V (Pacific Blue™, RRID: AB_1279044) and 7-AAD (BioLegend, Cat. No. #420404) to the manufacturer's instructions. Samples were acquired on a BD LSRFortessa equipped with a high-throughput samples (HTS) (BD Biosciences).

Adult AML patient samples were obtained from the Quebec Leukemia Cell Bank (BCLQ) under approved Research Ethics Board (REB) protocols and cultured in OCI-AML-8227 media supplemented with 500 nM of SR1 (Cat. No.: #72342) and UM729 (Cat. No.: #72332) (STEMCELL Technologies). Cells were treated under the same conditions as above. Cells were stained with True-Stain Monocyte Blocker™ (Cat. No.: #426102) and antibodies to CD45 (AlexaFluor®700, RRID:AB_2566372), CD19 (BV711, RRID:AB_2562065), CD33 (APC, RRID:AB_314352), CD11b (BV650, RRID:AB_2563793), CD15 (FITC, RRID:AB_314196), CD34 (APC-Cy7, RRID:AB_1877168), and CD38 (PE, RRID:AB_2561900) (BioLegend), followed by Annexin V (Pacific Blue™) and 7-AAD. Flow cytometry was performed as above. Sex and age of patient samples were not considered as a biological variable.

**Colony forming unit assay**

Colony forming unit (CFU) assays were performed to evaluate the functional impact of candidate compounds on progenitor activity. Drug concentrations were selected based on doses previously tested, using the next two highest concentrations above the $LC_{50}$ of $CD34^{+}CD38^{+}$ cells for each compound in both CB and OCI-AML-8227. Triplicate wells were pooled, washed with 1X PBS, and viable cells from the DMSO control were counted to establish a reference. Equal volumes of cell suspension were used across all conditions, corresponding to 8 000 cells for OCI-AML-8227 and 500 cells for cord blood. Cells were diluted in Iscove's Modified Dulbecco's Medium (IMDM) (Life Technologies) with 2% fetal bovine serum (Wisent) and plated in duplicate in MethoCult™ H4435 Enriched medium (#04435, STEMCELL Technologies). Colonies were assessed after 12 days, with colony types identified by morphology and color using standard criteria and quantified relative to DMSO control plates.

**Toxicity of top candidates on $CD34^+$ cord blood cells vs LSC-enriched OCI-AML-8227 cells**

Cord blood (CB) was obtained from healthy full-term deliveries under informed consent, following protocols approved by the REB of Héma-Québec, CHU Sainte-Justine, Université de Montréal, and the McGill University Health Centre. Mononuclear cells were isolated using Ficoll-Paque™ (GE Healthcare) and enriched for $CD34^+$ cells with the EasySep™ CD34 Positive Selection Kit (STEMCELL Technologies). Sex was not a biological factor. $CD34^+$ CB cells were cultured in SFEM II (STEMCELL Technologies) supplemented with 1% penicillin-streptomycin (Life Technologies), 10 ng/mL IL-6, 100 ng/mL SCF, 100 ng/mL FLT3L, 10 ng/mL G-CSF, 15 ng/mL TPO (Life Technologies), 500 nM SR1 and UM171 (STEMCELL Technologies). Cells were treated with four top candidate compounds at six doses (0.02% DMSO, 1.95, 7.8, 31, 125, 500, and 2 000 nM) for six days. In parallel, OCI-AML-8227 cells (50 000 cells per well) were treated identically in corresponding media. Following treatment, cells were stained for flow cytometry using antibodies CD34 (APC, RRID:AB_1877153), CD38 (PE, RRID:AB_2561900), CD15 (FITC, RRID:AB_314196), and CD45 (AlexaFluor®700, RRID:AB_2566372) (BioLegend) (OCI-AML-8227); CD7 (PE-Cy7, RRID: AB_2563941), CD10 (PE-Cy5, RRID: AB_314917), CD34 (APC-Cy7, RRID:AB_1877168), CD38 (PE, RRID:AB_2561900), CD45 (AlexaFluor®700, RRID:AB_2566372), CD45RA (BV650, RRID: AB_2563653), CD49f (APC, RRID: AB_1575047), CD90 (PE/Dazzle™, RRID: AB_2566343), and CD135 (PerCP-Cy5.5, RRID: AB_2565548) (BioLegend) (for CB). SYTOX™ Blue Dead Cell Stain (Life Technologies) was used to assess viability. Flow cytometry was performed using a BD LSRFortessa equipped HTS (BD Biosciences).

**Intracellular staining with cell surface staining**

Cells were collected and washed with PBS (pH 7.4, Gibco™, Fisher Scientific), then stained with BD Horizon™ Fixable Viability Stain 575 (1:1000; BD Biosciences, Cat. No.: ) for 15 minutes at room temperature to exclude dead cells. Surface staining was performed by incubating cells with CD34 (APC) and CD38 (PE) (BioLegend) in 2% CCS in PBS (Fisher Scientific) for 45 minutes on ice in the dark. Cells were then washed and fixed/permeabilized using the BD Cytofix/Cytoperm™ Fixation/Permeabilization Kit (Cat. No.: #554714, BD Biosciences) following the manufacturer's protocol. Intracellular staining was performed using antibodies USP1 (AlexaFluor®488, Novus Cat# NB100-88117AF488, RRID:AB_3174053) (Novus Biologicals Canada) for 30 minutes at 4°C in the dark. Samples were acquired on an LSRFortessa (BD Biosciences) equipped with a HTS and analyzed using FlowJo™ software (v10).

#### *Ex vivo* treatment with candidate compounds

Mouse experiments were performed according to protocols approved by McGill University and its Affiliated Hospital's Research Institutes. Sex and age were considered biological variables. Xenotransplantation experiments were performed exclusively in female recipient mice of 8-12 weeks old. Subjects were grouped randomly and blinding of investigators during the conduct and analysis of the study was performed. Group size was based on previously approved protocols mentioned. Subjects were not included if did not reach endpoint.

OCI-AML-8227 cells were treated with 500 nM of BIO-acetoxime (APExBIO, Cat. No.: #B5488), SJB2-043 (APExBIO, Cat. No.: #A3823), UMxxxxx03 (IRIC), or 0.05% DMSO (Fisher Scientific) as control. After 7 days, cells were collected, washed, and stained with True-Stain Monocyte Blocker™ (BioLegend, Cat. No. #426102), CD34 (APC, RRID:AB_1877153), CD38 (PE, RRID:AB_2561900) (BioLegend) and SYTOX™ Blue Dead Cell Stain (Pacific Blue™)

(Life Technologies) and analyzed on an LSRFortessa flow cytometer with high-throughput sampler (BD Biosciences).

For xenograft assays, NOD-SCID IL2Rg^null-3/GM/SF (NSG-S) female mice (Jackson Laboratories, RRID:IMSR_JAX:013062) were irradiated (2.1 Gy, X-RAD SmART Irradiator, Precision X-Ray, Inc.) 24 hours prior to intrafemoral injection of 25 μL of total OCI-AML-8227 cells (containing 1 000 $CD34^{+}CD38^{-}$ cells per DMSO control, $n = 5$ mice per condition). Twelve weeks post-injection, mice were sacrificed and bone marrow (injected and contralateral femurs) and spleen were collected. Human engraftment ($CD45^{+}$) and phenotype were assessed by flow cytometry using previously described antibodies (BioLegend).

**Statistics and reproducibility**

Data were expressed as means ± s.d. and statistical analysis was performed using GraphPad Prism v10 for [Windows/Mac] (GraphPad Software, Boston, MA, USA, RRID: SCR_002798) unless otherwise mentioned. $LC_{50}$ values were calculated using nonlinear regression (dose–response inhibition) in GraphPad Prism v10. For apoptosis a cut-off of 5% total apoptosis (total Annexin V+) was used throughout the study. When two-group comparisons were performed, a two-sample unpaired Student's *t* test was used. For multiple group comparisons, one-way ANOVA followed by Dunnett's multiple comparison was used. These tests were performed using GraphPad Prism v10 for [Windows/Mac]. For primary hit selection and normalization, statistical analysis is explained in Supp. Methods. Volcano plots used a cut-off of $|log_2FC>1|$ and FDR < 0.001 for significant genes. Statistical significance of ssGSEA enrichment scores was calculated using a two-sided Wilcoxon test. $P < 0.05$ was considered statistically significant. In the figures, asterisks indicate *$p < 0.05$, **$p < 0.01$, ***$p < 0.001$ and **** $p < 0.0001$ and "ns" being non-significant.

Flow cytometry data was analyzed using Flowjo™ Software Version 10.10.0 (Ashland, OR: Becton, Dickinson and Company; 2023).

**Single-cell RNA sequencing**

OCI-AML-8227 cells were enriched using SFEM II and the EasySep™ Magnet and EasySep™ CD34 Positive Selection Kit II protocol (Catalog #17856, STEMCELL Technologies). A total of four washes were performed instead of five to keep some $CD34^-$ blasts in the purified sample. Cells were then washed in SFEM II and counted using a hemocytometer to determine final yield. Some cells were stained with antibodies against CD34 (APC, RRID:AB_1877153), CD38 (PE, RRID:AB_2561900), CD15 (FITC, RRID:AB_314196) (Biolegend) and SYTOX™ Blue Dead Cell Stain (Life Technologies). Flow cytometry was performed using a LSR Fortessa to determine purity of $CD34^+$ and $CD34^-$ fractions. Cells were then plated (50,000 cells/well) and treated with 0.05% DMSO only (Fischer Scientific), 500 nM BIO-acetoxime, 250 nM SJB2-043 and 1 000 nM UMxxxxx03.

Cells were incubated for 24 hours and then collected, washed and resuspended in PBS with 0.04% BSA (Life Technologies). We aimed to capture ~10 000 cells per sample. An aliquot of cells was used for LIVE/DEAD viability testing (Thermo Fisher Scientific). Single-cell libraries were generated using the 10x Genomics Chromium X instrument and Chromium Next GEM Single Cell 3′ GEM, Library & Gel Bead Kit v3.1 (10x Genomics) according to the manufacturer’s protocol. Briefly, cells suspended in reverse transcription reagents, along with gel beads, were segregated into aqueous nanoliter-scale gel bead-inemulsions (GEMs). The GEMs were then reverse transcribed in a T1000 Thermal cycler (Bio-Rad) programed at 53ºC for 45 min, 85ºC for 5 min, and hold at 4ºC. After reverse transcription, single-cell droplets were broken, and the single-strand cDNA was isolated and cleaned with Cleanup Mix containing DynaBeads (Thermo Fisher

Scientific). cDNA was then amplified with a T1000 Thermal cycler programed at 98ºC for 3 min, 12 cycles of (98ºC for 15 s, 63ºC for 20 s, 72ºC for 1 min), 72ºC for 1 min, and hold at 4ºC. Subsequently, the amplified cDNA was fragmented, end repaired, A-tailed and index adaptor ligated, with SPRIselect Reagent Kit (Beckman Coulter) with cleanup in between steps. Post-ligation product was amplified with a T1000 Thermal cycler programed at 98ºC for 45 s, 12 cycles of (98ºC for 20 s, 54ºC for 30 s, 72ºC for 20 s), 72ºC for 1 min, and hold at 4ºC. The sequencing-ready libraries were cleaned up with SPRIselect, quality controlled for size distribution and yield (LabChip GX Perkin Elmer), and quantified using qPCR (KAPA Biosystems Library Quantification Kit for Illumina platforms).

Libraries were loaded on NovaSeq 6000 and sequenced using the following parameters: 28 bp Read1, 8 bp Index i7, 8 bp Index i5 and 150 bp Read2.

Raw sequencing BCL files from the Illumina sequencing were demultiplexed into paired-end, gzip-compressed FASTQ files using Illumina's bcl2fastq. Following this using cellranger count version 7.1.0 (10X genomics), reads were aligned to the GRCh38 human reference genome, and transcript counts were quantified for each annotated gene within every cell. The resulting UMI count matrices (genes Å~ cells) were then provided as input to Seurat suite.

**Single-cell RNA sequencing analysis**

Cell Ranger v7.1.0 (10x Genomics) was used to map reads to the human reference transcriptome, GRCh38 v1.2.0 (also from 10x Genomics). Seurat v5.1.0 (2) within R v4.3 (3) for was used for data normalization, integration and visualization. Cells were filtered out if they had: more than 15% mitochondrial gene expression, less than 250 genes identified, less than 500 UMIs as well as, those cell that had a low novelty score (log10 GenesPerUMI < 0.8). Similarly, the described steps by DoubletFinder v2.0.4 (RRID:SCR_018771) (4) were followed to identify and remove doublets from the dataset.

"SCTranform" was used to perform normalization regressing out mitochondrial expression, followed by integration as described by Seurat using the "RPCA" method. The umaps, violin plots, heatmaps and related figures were made using a combination of Seurat v5.1.0 (RRID:SCR_007322) and Tidyverse v2.0.0 (5) tools.

**Cell type prediction using ANNCAST**

Cell type prediction was performed using the ANNCAST classifier v1.2 (6), an artificial neural network-based model trained specifically for AML cells. To ensure compatibility with the classifier, outdated gene symbols from Ensembl84 were first converted to Ensembl97 by mapping GeneIds to the appropriate gene symbol version used by ANNCAST. Gene expression matrices were then normalized by sequencing depth, scaled to a constant depth of 10 000 UMIs, and log-transformed. To reduce the likelihood of ambiguous cell type assignments, any cell with a confidence score below 0.8 was re-assigned to the majority cell type among its five nearest neighbors (based on PCA embeddings) that had high-confidence scores (≥0.8). Because the ANNCAST classifier assigns cells to one of 18 high-confidence subtypes—and some of these subtypes contained relatively few cells, these were further grouped into seven broader categories: LSC, MultiLin-GMP, LMPP, Promyelocyte, S100A+ preNeutrophil, and Monocyte/MoDC/Mac, Erythrocyte/Megakaryocyte/Mast progenitors were also grouped to obtain subsets of suitable size for differential gene expression analysis. The resulting cell type assignments were reviewed and validated using a panel of key cell lineage marker genes.

**Differential gene expression (DGE)**

Differential gene expression (DGE) analysis was performed to assess the impact of treatment on the transcriptional profile. Each treatment condition was compared to the DMSO control. Comparisons were conducted within unsupervised clusters, within ANNCAST-categorized cell types, and within cell types stratified by tumor subclone identity. A zero-inflated model was applied to the log-normalized gene expression data using the MAST v1.28 (RRID:SCR_016340) statistical framework(7). The number of detected genes per cell was included as a covariate alongside the treatment effect. P-values were adjusted for multiple testing using the Benjamini-Hochberg (FDR) method. For the DGE analysis within tumor subclones, only clone–cell type combinations with at least 100 cells were included in the analysis to ensure robust differential expression testing.

### References

1. Safa-Tahar-Henni S, Páez Martinez K, Gress V, Esparza N, Roques É, Bonnet-Magnaval F, et al. Comparative small molecule screening of primary human acute leukemias, engineered human leukemia and leukemia cell lines. Leukemia. 2025;39(1):29-41.

2. Hao Y, Hao S, Andersen-Nissen E, Mauck WM, 3rd, Zheng S, Butler A, et al. Integrated analysis of multimodal single-cell data. Cell. 2021;184(13):3573-87.e29.

3. Team RC. R: A language and environment for statiscal computing. Vienna, Austria: R Foundation for Statistical Computing; 2021 [Available from: https://www.R-project.org/.

4. McGinnis CS, Murrow LM, Gartner ZJ. DoubletFinder: Doublet Detection in Single-Cell RNA Sequencing Data Using Artificial Nearest Neighbors. Cell Systems. 2019;8(4):329-37.e4.

5. Wickham H, Averick M, Bryan J, Chang W, D'Agostino McGowan L, François R, et al. Welcome to the tidyverse. Journal of Open Source Software. 2019;4(43):1686.

6. Khakipoor B, Lesi V, Farah A, Gingras O, Lavallee V-P. ANN-CAST : Automated Neural Network Classification for AML Single-Cell Transcriptomes. 2024.

7. Finak G, McDavid A, Yajima M, Deng J, Gersuk V, Shalek AK, et al. MAST: a flexible statistical framework for assessing transcriptional changes and characterizing heterogeneity in single-cell RNA sequencing data. Genome Biol. 2015;16:278.
